# Systematic functional drug testing in patient-derived models reveals ex vivo sensitivities associated with clinical outcome in rare solid tumors

**DOI:** 10.64898/2026.02.18.705724

**Authors:** Jasmina Paluncic, Zunamys I. Carrero, Lilly K. S. Fischer, Juliana P. Schulz, Dorothea Hanf, Attila Jády, Fatima Gutierrez Tenorio, Anna Klimova, Claudia Dagostino, Isabell Wolf, Melanie Hüther, Maximilian Werner, Parham Pourabbas Tahvildari, Sara Hrabovska, Maria Gabriela Pereira dos Santos, Elinor Young, Ivona Mateska, Sabine Schulze, Rebecca Prause, Heike Peterziel, Johanna Kirchberg, Daniel E. Stange, Benjamin Schmidt, Daniel Huebschmann, Claudia Scholl, Martin Schneider, Dana Westphal, Alexander A. Wurm, Ina Oehme, Olaf Witt, Jessica Pablik, Vivek Venkataramani, Christoph E. Heilig, Simon Kreutzfeldt, Peter Horak, Lino Möhrmann, Irina A. Kerle, Stephan M. Richter, Jürgen Weitz, Klaus-Dieter Schaser, Daniela Richter, Stefan Froehling, Christoph Heining, Hanno Glimm, Claudia R. Ball

## Abstract

Rare cancers are individually uncommon but collectively represent a substantial share of cancer burden, with limited systemic treatment options for many entities. Molecular profiling identifies targetable alterations, but actionable findings are limited and responses can vary despite a matched target. This motivates complementary approaches that directly assess tumor drug response. Here, we establish a biopsy-compatible *ex vivo* drug sensitivity testing platform optimized for low input and reproducibility. Patient-derived material was tested either directly or following *ex vivo* expansion. Functional profiling was performed within clinically relevant timelines across models from 126 patients with rare advanced solid tumors. Drug responses were consistent between model types. In most samples, we identified at least one potentially active compound, supporting feasibility at biopsy-scale. High *in vitro* sensitivity was associated with clinical benefit and progression-free survival. These findings support functional drug sensitivity testing as a complementary component in precision oncology for adults with rare cancers.

**Statement of Significance:** This study presents a biopsy-compatible drug sensitivity testing platform for phenotype-based therapy stratification in rare cancers. It identifies actionable ex vivo drug responses and shows associations with clinical outcome in patients treated with screened therapies. These findings support functional testing as a complementary additional layer of stratification for therapeutic prioritization.

## Introduction

Rare cancers account for approximately 20% of all malignancies, but mostly have very limited systemic treatment options, and clinical outcomes are frequently poor (1–3). Although systemic treatment options are available, selecting the most appropriate therapy remains challenging, particularly in the context of molecular heterogeneity and disease progression. The low incidence of individual entities limits prospective clinical trials, and the large number of distinct molecular and clinical subtypes results in few well-established, evidence-based treatment strategies, especially after failure of standard therapies (1–4). Molecular profiling has become a key tool to identify targetable alterations across cancer types, including NTRK and ALK fusions, FGFR2/3 alterations, IDH1 mutations and BRAF-driven tumors (5–8). However, such biomarkers are identified in only a subset of patients, and clinical responses vary even when a targetable alteration is present (7,9,10). This underscores the need to complement genomic and transcriptomic analyses with strategies that help to identify molecularly defined responders at the phenotype level. Ex vivo drug sensitivity testing offers such a complementary approach by directly measuring drug response in patient tumor cells (10). It adds a functional readout that can capture context-dependent cell states and treatment sensitivities that cannot be evidenced from sequencing alone (11–14). Proof-of-concept studies in solid tumors have shown that functional testing can identify clinically relevant vulnerabilities in defined settings, including MEK inhibitor sensitivity in MAPK-driven pediatric tumors, PARP inhibitor response in DNA repair-deficient models or kinase inhibitor susceptibilities in individual cases (14–17). However, whether such approaches are feasible and informative for adults with rare solid tumors in routine, biopsy-based settings remain unclear. Key challenges include limited access to viable tissue, biological heterogeneity, and lack of standardized workflows, which complicate reproducibility and clinical interpretation. Consequently, there is a need for standardized, biopsy-compatible functional testing strategies that integrate with molecular profiling and can be applied across diverse rare solid tumors with consistent assay performance and clinically meaningful readouts.

In this study, we establish and apply a biopsy-compatible ex vivo drug sensitivity testing (DST) strategy for advanced rare solid tumors. We present a scalable framework for profiling of clinically relevant anticancer agents across short- and long-term patient-derived models, assess assay reproducibility and evaluate associations between functional readouts and clinical outcomes in patients. Together, these data provide a framework for integrating functional testing with molecular profiling in precision oncology for adults with rare cancers, a population with substantial unmet clinical need.

## Results

### Establishment of a miniaturized *ex vivo* drug sensitivity testing platform

Functional drug sensitivity testing can complement molecular profiling for patients with rare malignancies, but its broader use is constrained by assay complexity and limited biopsy material. We therefore established a miniaturized *ex vivo* drug sensitivity testing (DST) platform optimized for low cell input, technical reproducibility and compatibility with automated pre-spotted plates **(Fig. 1A, Fig. S1).** To enable a cell-sparing assay while preserving reliable dose-response behavior, we evaluated whether reproducible high-resolution dose-response curves over 20 concentrations **(Fig. S1A)** could be reliably compressed. Using representative patient-derived models, *in silico* and experimental downscaling demonstrated that 10-, 5- and 3-point concentration series retained characteristic sigmoidal curves and resulted in comparable IC50 estimates relative to 20-point curves **(Fig. 1B).** Based on this, we selected a 5-point concentration design to substantially reduce cell numbers while maintaining reliable IC50 estimation. We next assessed the cell input effect on DST performance. Dose-response curve quality and IC50 values remained consistent over a wide range of seeding densities **(Fig. S1B-C),** with curve fitting and IC50 estimation being maintained down to 100 cells per well. A seeding density of 500 cells per well was selected as a robust lower bound for subsequent experiments.

**Figure 1.**
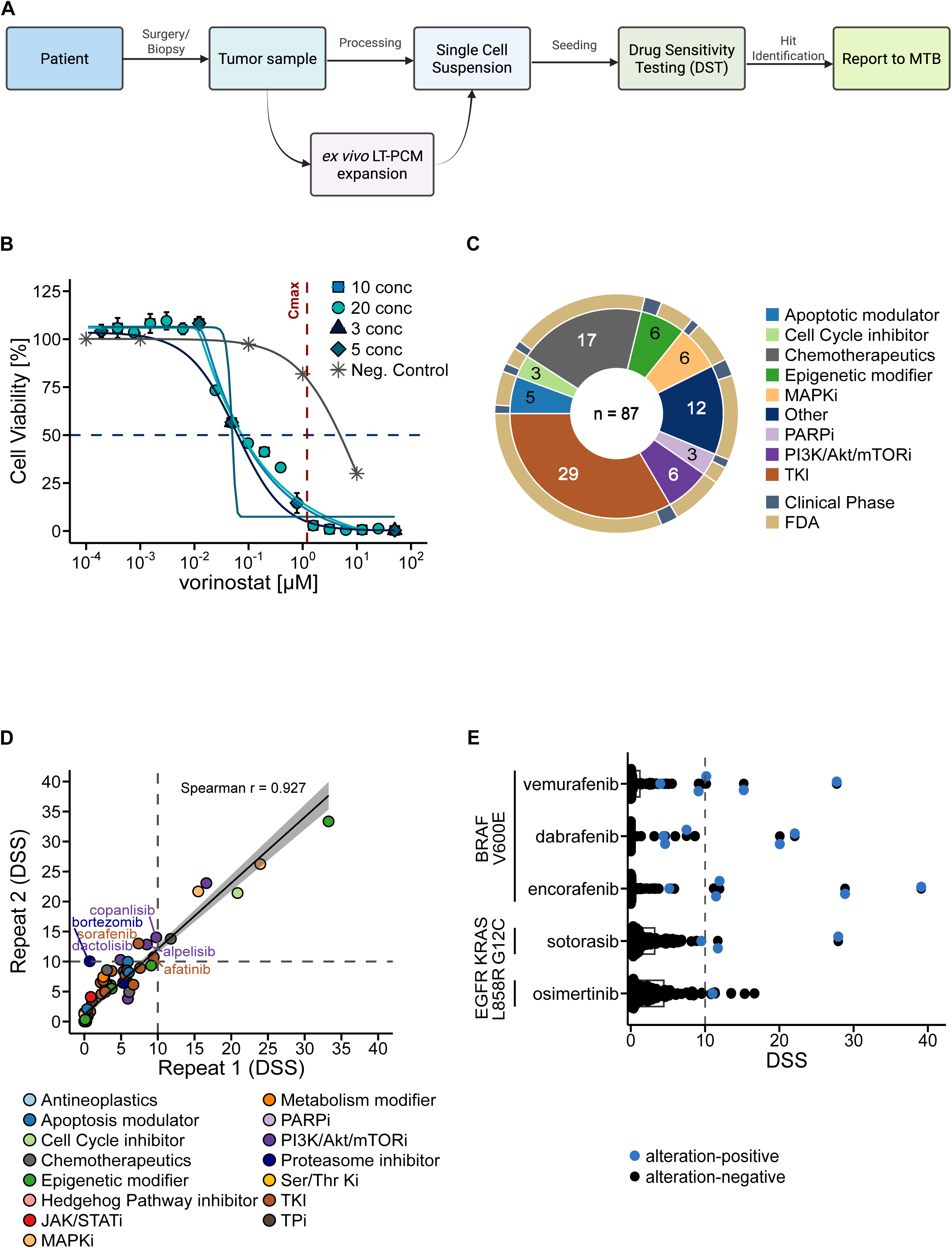
Miniaturized ex vivo Drug Sensitivity Testing (DST) enables robust, low-input functional profiling of patient-derived tumor models. **A** Schematic overview of the miniaturized DST workflow. Freshly dissociated tumor samples or pre-expanded PCMs are seeded as short-term cultures into 384-well plates and exposed to a 5-point concentration series of an 87-compound drug panel. Cell viability is quantified after 72 hours using ATP-based luminescence, and drug sensitivity is summarized using the DSS metric (DSS_asym_adj_). Identified ex vivo active substances (Hits) are then reported in the molecular tumor board (MTB). **B** In silico and experimental downscaling of high-resolution dose-response curves containing of 20-concentrations demonstrates that 10-, 5-, and 3-concentration series retain characteristic sigmoidal profiles and produce IC50 values comparable to the original 20 concentration format. Shown are representative examples from patient-derived models. These results support the selection of a 5-point format as a cell-sparing yet quantitatively stable assay design (n=2 biological replicates). **C** Overview of the clinically relevant 87-compound library enriched for targeted agents applicable in biomarker-driven therapy decisions in rare and molecularly heterogeneous cancers. The panel summarizes drug class composition and clinical development status of the library. **D** Concordance of drug response profiles between manually prepared dilution plates and automated pre-spotted plates. Exemplary dose-response curves and DSS comparisons show strong reproducibility across plate formats and across biological replicates, confirming robustness of the miniaturized high-throughput screening configuration (n=1 biological replicates). **E** Mutation-specific drug responses validate pharmacological specificity of the platform. Genotype-matched models (BRAFV600E, KRASG12C, EGFRL858R) show higher DSS values and enrichment of responders to corresponding targeted inhibitors (encorafenib n=5, dabrafenib n=5, vemurafenib n=5, sotorasib n=3, Osimertinib n=1), whereas mutation-negative controls show generally lower DSS values (encorafenib n=109, dabrafenib n=109, vemurafenib n=115, sotorasib n=121, osimertinib n=113). Dose-response curves and DSS values illustrate the preservation of on-target effects under reduced-input, miniaturized testing conditions.

To validate platform performance under screening conditions, we used pre-spotted 384-well plates containing a clinically relevant 87-compound panel enriched for targeted agents applicable in rare and molecularly heterogeneous cancers **(Fig. 1C)**. Drug sensitivity was quantified using the adjusted asymmetric Drug Sensitivity Score (DSS_asym,adj;_ referred to as DSS for simplicity) (17,18). A DSS ≥10 was used to define functionally active compounds in patient-derived cells, defined by a measurable *ex vivo* drug response in tumor cells (“Hit”) (18). Dose-response profiles, DSS values and hit classifications showed strong concordance between manually prepared plates and pre-spotted plates and between biological replicates **(Fig. 1D, Fig. S1D)**. Readout was stable following plate storage under standardized conditions for up to 27 weeks **(Fig. S1E-F)**, supporting advance preparation of pre-spotted plates without affecting readouts. To assess accuracy, we tested alteration-specific inhibitors in a set of engineered and patient-derived samples. Cells that carry an inhibitor-directed mutation showed DSS values ≥10 and were enriched among responders to corresponding targeted drugs, whereas mutation-negative controls showed generally lower DSS values (DSS<10) **(Fig. 1E**). Together, these data demonstrate that the miniaturized DST platform is technically robust and compatible with biopsy-level material.

### Drug sensitivity testing is reproducible across different PCM sources

We next examined whether drug sensitivity profiles remain consistent when functional testing is performed either directly on short-term patient-derive cancer models (ST-PCMs) or on PCM that were first expanded as long-term cultures (LT-PCMs). This comparison is practically relevant as biopsy material in rare-cancer settings frequently requires interim expansion, e.g. as organoids or spheroids, three-dimensional cell aggregates cultivated with or without Matrigel, respectively, to generate sufficient cell numbers for DST. ST-PCMs retain the freshly dissociated cellular composition of the tumor sample, whereas LT-PCMs represent tumor-cell enriched cultures obtained through *in vitro* expansion.

When comparing all ST- and LT-PCMs, the proportion of identified hits per sample was comparable and showed no significant difference between input model types (**Fig. 2A**). To assess whether broader sensitivity patterns vary between input model types, we compared DSS distributions for all drugs profiled in ST-versus LT-PCMs (**Fig. 2B**). DSS distributions were additionally stratified by proliferative activity.

**Figure 2.**
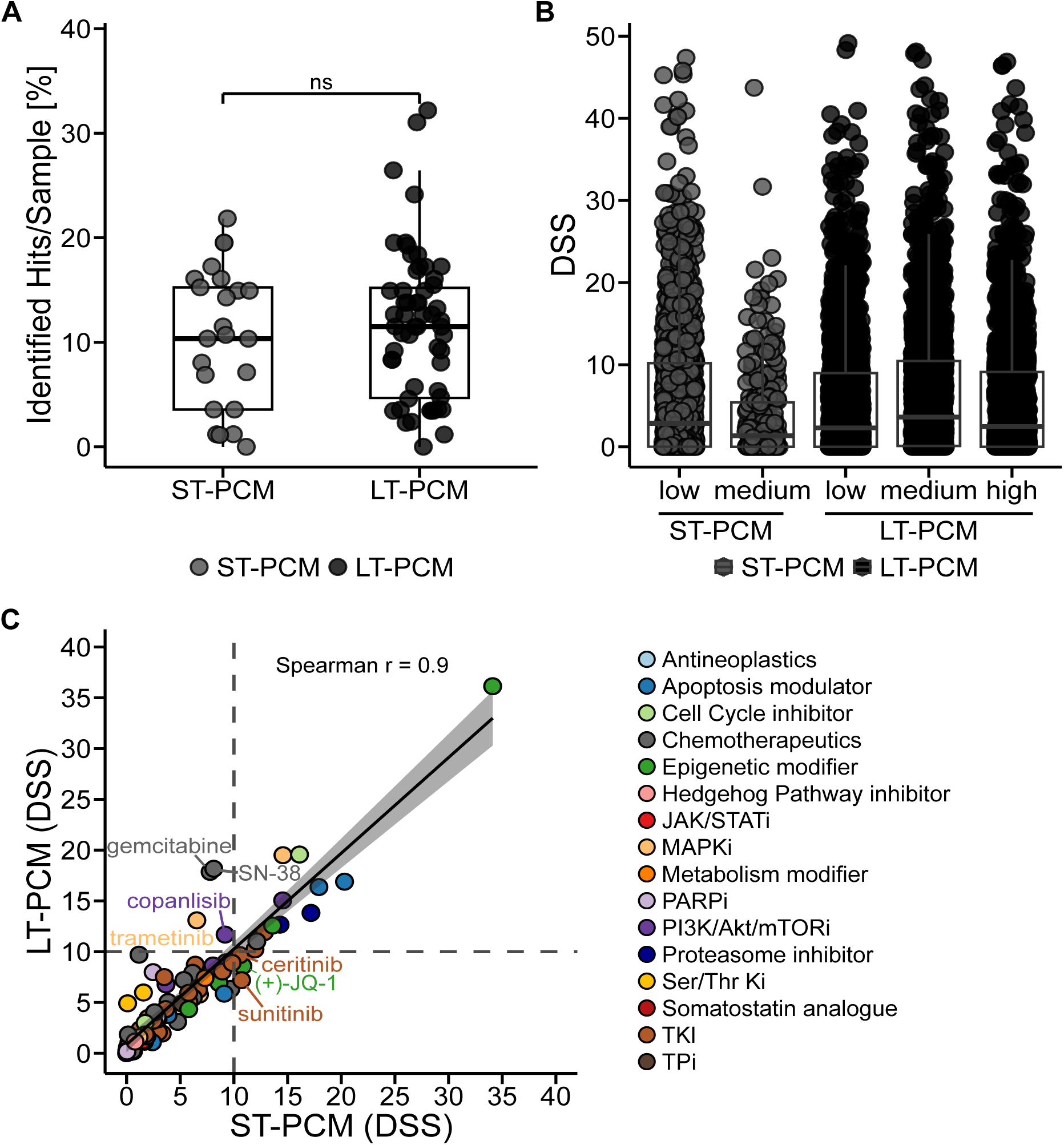
Drug sensitivity testing is reproducible across different PCM sources. **A** Distribution of hit counts per sample, normalized to the total number of drugs tested for that sample, between ST-PCM and LT-PCM. Samples are categorized as either ST-PCM with a minimum tumor cell content (TCC) of 80% (n=25) or LT-PCM (n=59). Boxes illustrate the 25th and 75th percentiles, respectively, while the center line represents the median. **B** Comparison of DSS distribution of samples treated with all 87 drugs. ST-PCM (n=18; TCC > 80%, full screen) or LT-PCM (n=53) were stratified based on their proliferation class, determined by an unsupervised k-means clustering approach, using the mean ATP readouts normalized to seeded cells per well as a surrogate of cellular proliferation. The center line represents the median and the whisker the standard deviation. Significance was determined by Mann-Whitney test (two-tailed). *p*-values< 0.05 were considered significant. **C** Averaged DSS Correlation between LT-(n=53) and ST-PCMs (n=25; TCC > 80%) treated with 84 drugs. For each drug, DSS values were averaged across LT-PCMs and ST-PCMs, and the resulting per-drug mean DSS values were correlated. The black diagonal represents the correlation line starting at 0. Colored labels indicate the drug class of the specific drug. Spearman’s rank correlation coefficient (r) is displayed.

The drug-specific average DSS values were highly similar and showed strong correlation between ST- and LT-PCMs **(Fig. 2C**), indicating that the general pattern of drug responses is preserved regardless of whether DST is performed on freshly dissociated short-term cultures or on long-term expanded models. This observation was further supported by drug class analysis, which showed broadly comparable DSS distributions between ST- and LT-PCMs across the major classes tested, and indicating some class-specific and proliferation-associated shifts in sensitivity (**Suppl. Fig. 2)**.

To assess model-specific concordance at the level of individual patients, we next evaluated cases where different PCMs or PCM types were available from the same tumor specimen. Comparable to biological replicates from the same sample, matched comparisons of different PCM from the same patient showed a strong correlation of DSS values without systematic gain or loss of drug sensitivity (**Fig. 1D, Fig. S1G-H**), indicating that patient-specific response patterns are retained in different PCM types. Comparable consistency was observed across common clinical handling conditions, including fresh versus frozen-thawed material and immediate versus short-term acclimatized processing, each showing strong correlation between DSS values (**Fig. S1I-J**).

Together, these results demonstrate that DST performance is consistent in heterogeneous PCM and under varying sample-handling conditions, supporting its feasibility for clinical and translational settings.

### Drug sensitivity testing is feasible in diverse PCM types and entities

We next applied *ex vivo* DST to clinically derived samples from patients with advanced rare solid cancers, where tumor material is frequently scarce and therapeutic options are limited **(Fig. 3)**. In total, we tested 137 PCMs generated from 126 rare advanced cancer patients from a precision oncology patient cohort within clinically relevant timelines **(Fig. 3A-C)**. Comprehensive genomic profiling confirmed the expected molecular heterogeneity of rare cancers, including recurrent alterations affecting TP53, CDKN2A, KRAS and additional oncogenic pathways **(Suppl. Fig. 3).** To provide broader biological context, 26 additional PCMs from our in-house repository, representing both rare and more common tumor types, were included. Altogether, the full dataset comprised 163 patient-derived models, complemented by nine well-characterized cell lines used as technical controls. Patient samples originated from patient tumors or were pre-expanded as LT-PCMs (organoids, spheroids, semi-adherent cultures) or PDX, and all models were dissociated into single-cell suspensions prior to seeding, ensuring that DST was always performed on short-term culture-equivalent input material. Most tumor samples originated from biopsies (57.1%), including fine-needle punctures, while 42.9% originated from surgical resections.

**Figure 3.**
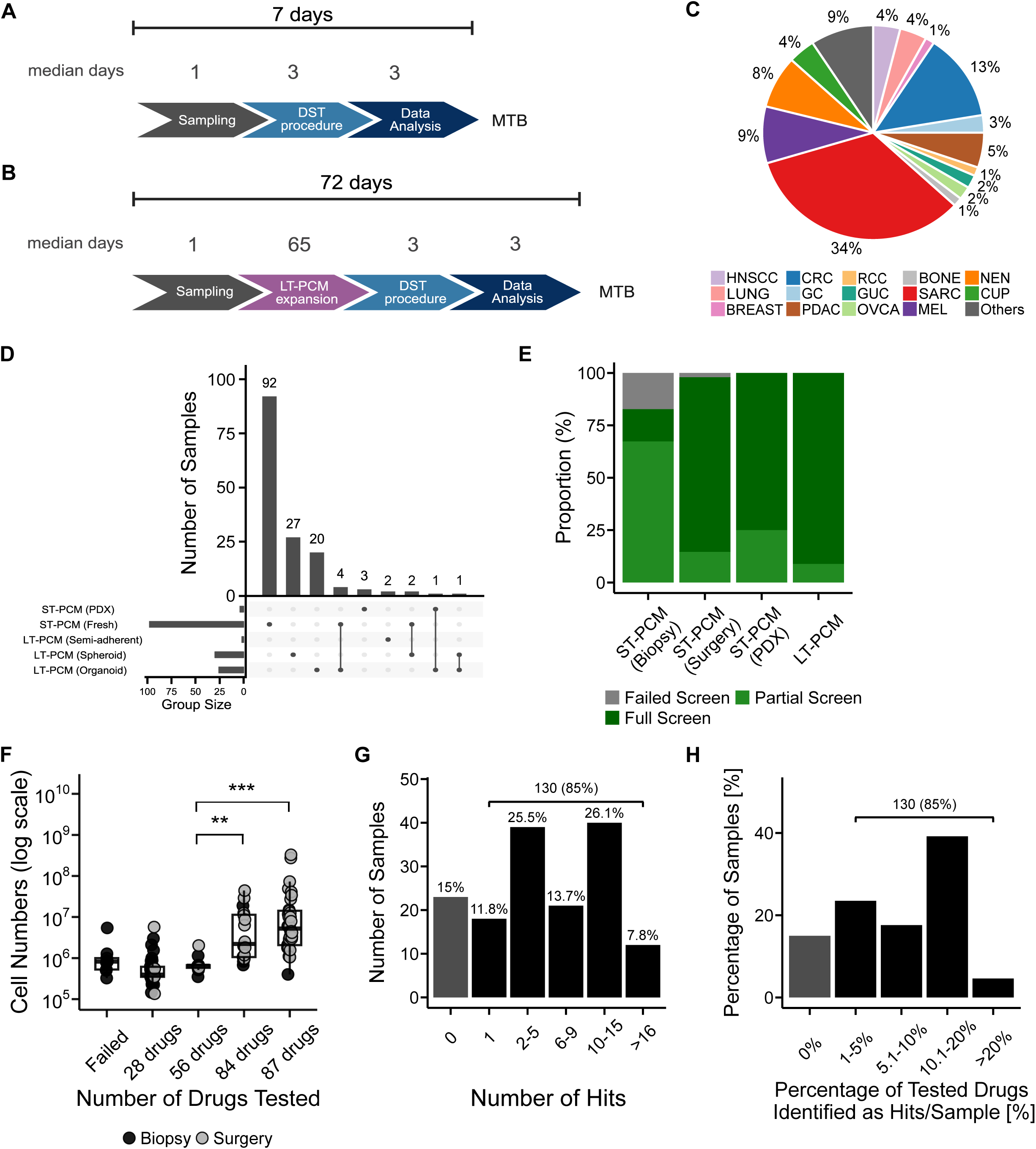
Drug Sensitivity Testing Captures Functional Responses Across Rare Solid Tumor Models. **A-B** Timeline from sample receipt to reporting of functional hits in **A** short-term (ST) and **B** long-term expanded (LT) PCMs. Numbers above the timelines represent median duration (days). **C** Tumor entity overview of all screened patient-derived tumor models (total: 28 entities, n=152). Entities occurring once were categorized as “other”. **D** UpSet plot illustrating intersections between different PCM types tested. Each vertical bar and black node (connected by a black line) indicate the number of patients from whom multiple PCM types were analyzed. Horizontal bars indicate the total number of patients within each individual PCM type group. ST-PCMs comprise short-term cultured cells derived directly from tumor tissue or following prior *in vivo* expansion in patient-derived xenografts (ST-PCM (PDX)). Spheroids and organoids are both long-term patient-derived cancer models (LT-PCMs) and are shown as distinct categories in the plot. **E** Proportion of failed, partial (28 or 56 drugs), or complete (84 or 87 drugs) drug screens by PCM sample of origin. **F** Cell count distribution of cell numbers grouped by the number of drugs tested per sample. Dots are colored according to the origin type. Significance was determined by the Wilcoxon Rank Sum test. **p<0.01, ***p<0.0001 **G** Number of hits per sample across various hit number categories, including the percentage of samples as well as total number and percentage of samples with a minimum of one hit. **H** Number of hits per sample, normalized to the total number of drugs tested per sample, shown across different hit rate groups. Bars indicate the percentage of samples in each hit-percentage bin, the bracket denotes samples which have at least one drug hit (>0%).

For patient samples from the precision oncology cohort (n=137), DST was successfully performed in 92.7% of cases (127/137). The median turnaround time from DST initiation to completion of functional testing was 7 days **(Fig. 3A).** For samples requiring ex vivo expansion prior to DST, the screening time from sample acquisition to DST result was driven by PCM generation and expansion (median 72 days) (**Fig. 3B)**. The combined PCM cohort (n=163) comprised patients with rare solid tumor entities with an annual incidence <6/100,000, including sarcomas (34%), as well as tumor types that have limited therapeutic options, including cancer of unknown primary (4%) and neuroendocrine tumors (8%) **(Fig. 3C).** A smaller subset represented more common solid tumors, several of which reflected early-onset disease or rare molecular subtypes within otherwise common entities.

In eight patients, multiple PCM types were tested, predominantly from the same patient specimen (n=7) and/or from different specimens of the same patient (e.g. distinct tumor locations (n=3) or longitudinal samples (n=1)). ST-PCM generation was prioritized whenever possible (63.8%, 104/163), while samples with insufficient cellular yield were pre-expanded as spheroids (18.4%), organoids (16.6%), PDX-derived short-term PCMs (2.5%), or semi-adherent cultures (1.2%) **(Fig. 3D)**. Among the subset of 126 patients with available clinical data, most had metastatic, treatment-refractory disease (51.6%) and had received multiple prior therapy lines **(Suppl. Table 1)**, reflecting the advanced disease setting in which functional testing was conducted.

### DST is feasible in clinical tumor samples despite limited cellular input

We next evaluated how clinical sample type (biopsy/surgical resection), cell input model (ST-PCM/LT-PCM/PDX) and input cell number affect feasibility and success of drug sensitivity testing in clinically relevant models **(Fig. 3E)**. Drug screens were classified as complete (≥3 plates with zprime_R≥0)), partial (1-2 plates with zprime_R≥0), or failed (zprime_R<0 on all plates). LT-PCMs and surgery-derived ST-PCMs consistently enabled complete or partial screening (97.2%) where zprime_R is a plate-quality metric based on the separation of positive and negative controls, with higher values indicating better assay performance. Biopsy-derived ST-PCM, which inherently provide fewer cells, showed greater variability, but DST remained feasible in the majority of biopsies that could be directly seeded on at least one drug plate (82.7%) **(Fig. 3E)**. Overall, screening success was strongly associated with total available cell numbers. Surgery-derived ST-PCM provided significantly higher median cell counts than biopsy-derived models (5.13×10⁶ vs. 6.25×10⁵; p-value=1.15×10⁻^9^) **(Suppl. Fig. 4A),** and increasing cell numbers were associated with a higher number of successfully tested drugs (median: 87 drugs=5.28×10^6^ cells; failed=8.25×10⁵ cells) (**Fig. 3F).** Functional profiling was achievable in 83% of seeded biopsy-derived models (44/53), demonstrating assay robustness even with minimal cellular input.

### High hit rates in patient-derived models demonstrate clinical feasibility of drug sensitivity testing

To evaluate the potential clinical relevance of drug sensitivity testing in rare and advanced cancers, we analyzed hit rates across 163 analyzable PCMs (n=152 patients; **Fig. 3G, Fig. S4B,).** At least one *ex vivo* hit was identified in 85% of cases, with a median of 5 hits per sample for the entire cohort and a median of 7 hits among samples in which at least one hit was detected. Most models showed moderate number of hit counts (1-15/sample), corresponding to approximately 1-20% of tested substances **(Fig. 3G-H**). Importantly, this approach was able to discriminate subgroups of responders and non-responders within drug classes **(Fig. S4C-F)**. Together, these findings indicate that patient-specific functional drug profiling can reveal distinct drug sensitivities in rare and advanced cancers.

### Entity- and pathway-associated drug response patterns

To determine whether functional drug responses reveal shared biological features among tumor models, we analyzed DSS profiles from 143 patient-derived PCMs using unsupervised clustering **(Fig. S5A)**. Drug response patterns were highly heterogeneous, and cultures rarely grouped by tumor entity or PCM type. Instead, clustering was mainly driven by pharmacological class, with compounds targeting PI3K, BRD, TOP2A, CDK4/6 and BRAF forming coherent activity blocks. This indicates that shared mechanism-of-action rather than tumor category accounts for the dominant structure in the dataset. Entity-associated trends were present, but less pronounced. MEK-pathway inhibitors showed general strong activity in multiple models, with CRC models displaying the highest sensitivity, consistent with frequent MAPK-pathway alterations in this entity (**Fig. S5B-D**). In contrast, sarcoma PCMs demonstrated more variable responses to both targeted and cytotoxic agents, reflecting the biological diversity characteristic of these tumors. Hit-frequency normalization confirmed these patterns and did not identify strong entity-wide deviations. Cytotoxic agents including gemcitabine, doxorubicin, trabectedin and etoposide exhibited activity in subsets of sarcoma PCMs but without uniform sensitivity, underscoring inter-tumor heterogeneity even within single entities. Bleomycin showed a modest enrichment in sarcoma compared to other tumor types, suggesting potential subtype-associated vulnerabilities.

### PCM-derived drug sensitivity correlates with clinical outcome

To evaluate whether *ex vivo* drug response is associated with patient outcome, we correlated DSS values from PCMs of 126 patients with clinical treatment responses **(Fig.4)**. Because therapies administered long after sampling may be confounded by clonal evolution or interim treatments, we focused the clinical correlation on a defined subset of patients who received a drug included in the DST panel as the next systemic therapy after tissue acquisition. Twenty patients met these criteria and were evaluable for correlation of PCM results with clinical outcome. In most cases, combination therapies were used, of whom one or two matched agents were tested *in vitro*. In PCM of 8/9 patients with documented tumor shrinkage (partial or mixed response), the administered drug showed a DSS≥10. In contrast, PCMs from patients with stable disease (n=5) as well as PCM from 6 of 7 patients with progressive disease showed DSS values <10 (**Fig. 4A**). Consistently, DSS values were significantly higher in responders (median DSS=13.7) compared to patients with stable disease (median DSS=0.9; p=0.019) and non-responders (median DSS=1.56; p=0.047) (**Fig. 4A**).

**Figure 4.**
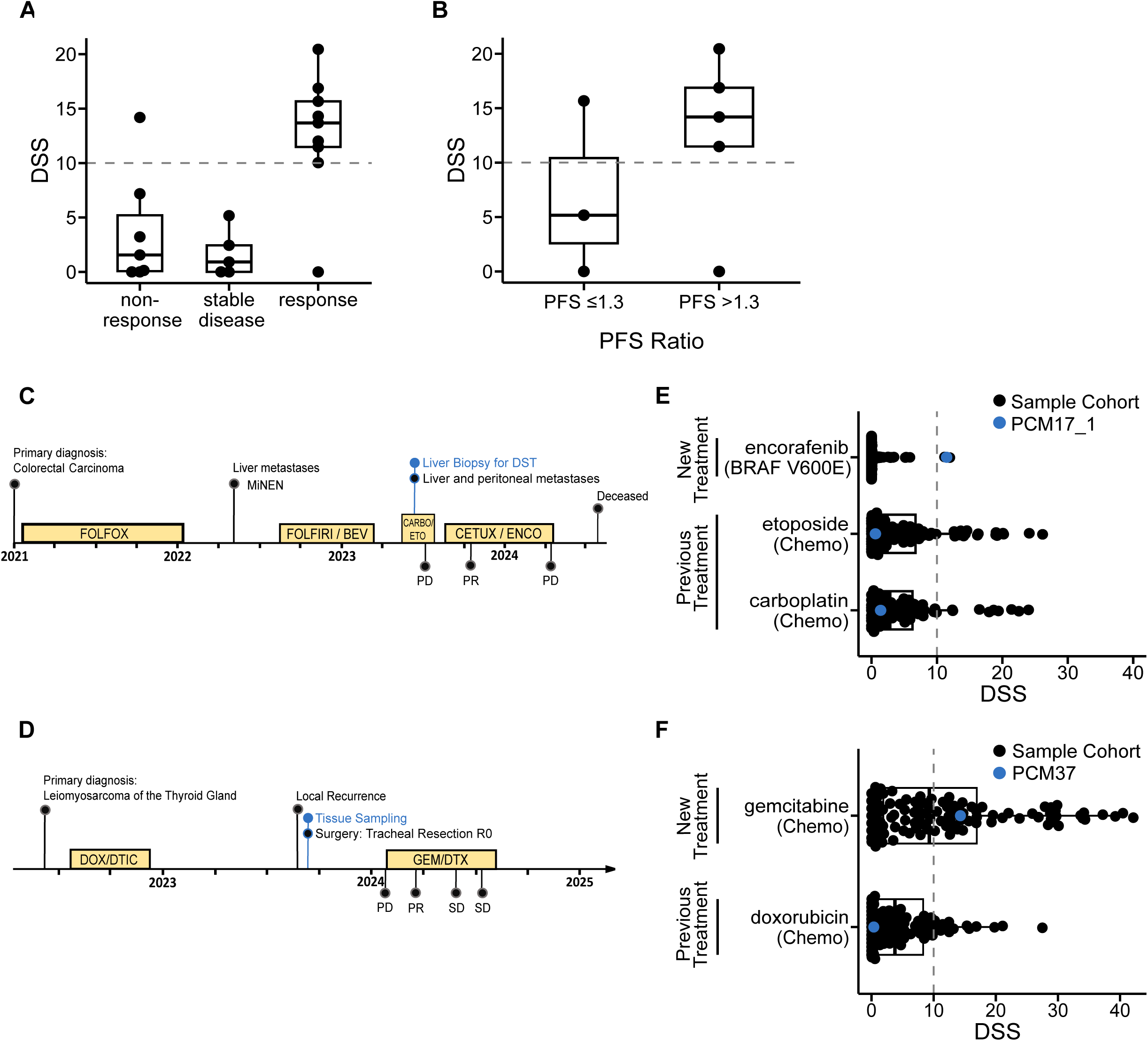
Predictive Capacity of in vitro Drug Response for Patient Outcome. **A-B** DSS distribution of samples treated with the patient’s received drugs as the next systemic therapy following sampling, categorized by **A** non-responders, patients with stable disease and responders or **B** ≤1.3 and ≥1.3 PFS Ratio. Patient response was defined as clear shrinkage of measurable tumor, but residual disease remains (partial response); simultaneous shrinkage of some lesions while others progress (mixed response); patient tumors with stable disease show no significant change. No response was defined as an increase in tumor size or the appearance of new metastatic lesions (progressive disease). Boxes illustrate the 25^th^- and 75^th^-percentiles, respectively, while the center line represents the median. Significance was determined *via* the Kruskal-Wallis test (Dunn Test with Bonferroni correction). **C** Patient example timeline including treatments, molecular alteration, *in vitro* drug sensitivity, and outcome of a patient diagnosed with colorectal carcinoma. **D** Patient example timeline including treatments, molecular alteration, *in vitro* drug sensitivity, and outcome of a patient diagnosed with leiomyosarcoma. **E** DSS distribution of all samples subjected to PCM17’s new and previous treatments (as monotherapies). DSS of PCM17’s PCM is indicated by the blue dot. **F** DSS distribution of all samples subjected to PCM37’s new and prior treatment, administered as monotherapies. DSS of PCM37’s PCM is indicated by the blue dot.

Progression-free survival ratio (PFSr) analysis, defined as the ratio of progression-free survival on the first treatment administered after tissue acquisition to that on the most recent prior therapy, was available for 8 patients and supported an association between higher DSS and clinical benefit. Patients with PFSr>1.3 had higher DSS values for the administered drug (median DSS=14.19) than those with PFSr<1.3 (median DSS=5.17; Kruskal Wallis rank-sum test, p-value = 0.02) (**Fig. 4B**). Finally, when including all treatment timepoints after tissue sampling where a PCM-tested drug was given to a patient (n=29 drugs from 20 patients), median DSS was again higher in responders (median DSS=13.7) compared to non-responders (median DSS=1.13, post-hoc Dunn’s test with Bonferroni correction, p=0.0056) and significantly higher compared to patients with stable disease (DSS=0.35, post-hoc Dunn’s test with Bonferroni correction, p=0.0082).

Two retrospective patients are highlighted to exemplify this association. A *BRAF^V600E^*-mutated CRC patient achieved 42-week partial response under cetuximab/encorafenib, with encorafenib ranking among the top 1% of DSS values in the corresponding PCM (PCM17, 99th percentile) (**Fig. 4C, E, Suppl. Table 1).** A leiomyosarcoma patient (PCM37) responded to gemcitabine/docetaxel for 25 weeks, with gemcitabine scoring DSS=14.3 (66th percentile), while doxorubicin showed minimal *in vitro* activity (DSS=0.37, 23rd percentile) (**Fig. 4D, F, Suppl. Table 1)**. In both cases, PCM-derived in vitro drug responses were consistent with observed clinical benefit.

In summary, PCM-derived drug sensitivity testing results were associated with observed clinical benefit across multiple analytical layers. A DSS≥10 distinguished responders from non-responders within this clinically defined cohort, supporting the potential of functional testing to inform individualized therapy selection in rare and advanced cancers.

## Discussion

To improve patient stratification for rare cancers, this study introduces a functional approach that refines patient stratification by integrating a miniaturized high-throughput drug sensitivity platform to identify actionable ex vivo therapeutic vulnerabilities, complementing sequencing-based target identification in rare cancer precision oncology. Across biologically and technically variable experimental conditions, this platform demonstrated robust and reproducible performance, supporting its applicability in settings with limited input material. Importantly, PCM-derived drug sensitivities were not only technically reproducible but also associated with clinical treatment outcomes, highlighting the potential of functional drug sensitivity testing to provide clinically relevant information beyond genomic profiling alone in rare cancer precision oncology.

Rare cancers remain a major challenge for precision oncology because patients often present with advanced disease, where surgical resection is not feasible and only diagnostic small biopsies are available for analysis. In such settings, limited cell yield restricts the feasibility of functional testing, while efficient protocols for generating expandable patient-derived models exist mainly for common entities such as colorectal, breast, and pancreatic cancers (19–21). For many rare tumors, however, model establishment remains inefficient or frequently fails, further limiting access to functionally testable material (22). A recent study showed the feasibility of short-term drug testing assays using freshly dissociated sarcoma specimens (14), but relied on large surgical samples and thus only partially reflects the clinical reality of scarce material from advanced or relapsed rare cancers (23). Our approach addresses this limitation by enabling DST directly on freshly dissociated tumor cells, thereby bypassing the need for long-term expansion. Using a miniaturized 384-well format, DST achieved reproducible results with minimal input and a turnaround time of approximately 7 days, which is compatible with clinical decision-making in advanced settings.

While success rates were highest for surgical specimen samples or LT-PCMs due to higher cell yields, drug testing could still be performed in biopsy-derived samples, indicating feasibility in clinically relevant specimen types. Nevertheless, low cell yield and limited tumor cell content of small biopsies remain major challenges for routine functional testing. The inclusion of long-term and PDX-derived PCMs can further expand assay coverage, enabling drug profiling in additional patient samples and thereby increasing insights into *ex vivo* drug vulnerabilities on a wide range of tumor entities.

Across the cohort, most samples responded to at least one drug *ex vivo*, regardless of sample type or origin. Minor differences in hit frequencies among PCM types from the same patient likely reflect intra-tumor heterogeneity and culture-associated effects. Long-term PCMs generally showed higher overall sensitivity than ST-PCMs, consistent with their greater proliferative activity, higher tumor cell content, and prior adaptation to *ex vivo* conditions. This increased responsiveness was most pronounced for compounds targeting proliferation, DNA damage, mitotic and cell cycle regulation. Such drugs may preferentially affect highly proliferative LT-PCMs, whereas freshly dissociated ST-PCMs might underestimate their efficacy due to lower proliferation and limited survival signaling after dissociation (24),(25). Despite these differences, drug response profiles were broadly concordant between model types, underscoring the robustness of our DST platform for functional testing in biologically distinct inputs.

In tumors with well-established therapeutic indications, PCM-derived drug hits matched the expected molecular targets, supporting assay specificity. Beyond these known associations, our cohort-based approach enabled relative comparison of drug sensitivities between entities, consistent with the INFORM study program (17). Notably, several patients who received DST-supported treatments achieved partial responses, underscoring the potential clinical relevance of functional profiling. Integrating functional data with multi-omics analysis may therefore help to identify therapeutic vulnerabilities in patients lacking established biomarkers or exhibiting acquired resistance in advanced or relapsed rare cancers. Consistent with previous precision oncology studies (11,12,17) our findings highlight the potential of DST to add functional information beyond genomic prediction, providing a complementary layer of information for patient stratification.

Our stringent quality and sensitivity criteria ensured high reproducibility and assay specificity. Importantly, mutation-specific inhibitors, such as those targeting EGFR, BRAF, or FGFR alterations, showed distinct activity patterns between responders and non-responders. This demonstrates that the applied thresholds reliably capture biologically and clinically relevant effects, including known drug-target relationships. For exploratory compounds or targets with less well-defined pharmacologic windows, a stepwise strategy combining broad primary screening with targeted follow-up validation could further enhance sensitivity while maintaining specificity - a principle similarly applied in other high-throughput screening settings such as CRISPR sgRNA library design and evaluation (26).

We identified *ex vivo* drug activities for advanced sarcoma- and CRC-derived models among compounds not conventionally used for those cancer types. Standard therapies less active in sarcoma PCMs, while TKIs and the HSP90 inhibitor ganetespib showed notable *ex vivo* activity, consistent with phase II trials reporting partial responses to sunitinib in alveolar soft part sarcoma and improved short-term progression-free survival with ganetespib in lipo- and leiomyosarcomas(27,28). In our cohort, PI3K inhibition also was more effective in sarcoma PCMs than in other tumor types, aligning with preclinical evidence from molecularly defined Ewing, synovial and alveolar rhabdomyosarcoma models (29). Further, combining standard chemotherapies with targeted agents, as demonstrated in the EORTC LMS-04 phase III trial where doxorubicin plus trabectedin prolonged progression-free survival compared to doxorubicin alone (30), supports the rationale for multi-agent functional testing (30,31). Collectively, these results highlight the potential of DST to nominate candidate drug sensitivities for further evaluation beyond established indications, while reproducing clinically observed sensitivity patterns. Our data further support that genomic and transcriptomic profiles alone may not fully capture functional dependencies, reinforcing the complementary role of functional testing. Considering additional regulatory layers, including post-translational modifications, may further refine biomarker discovery, especially in heterogeneous tumor cohorts. Expanding the PCM dataset could thus reveal novel, tumor-agnostic pathway dependencies that may inform individualized therapy development.

In conclusion, this study demonstrates that integrating DST into precision oncology is feasible and can reveal functional drug sensitivities beyond genomic prediction, with the potential to expand therapeutic options for patients with advanced and rare solid cancers. Its feasibility from limited input makes it particularly suited for clinical scenarios with scarce samples. Incorporating DST alongside genomic profiling within molecular tumor boards may enhance individualized treatment selection in a tumor-agnostic context and support more informed clinical decision-making.

## Methods

### Patient tissue

Fresh tumor tissue was obtained from patients enrolled in the DKFZ/NCT/DKTK Molecularly Aided Stratification for Tumor Eradication Research (MASTER) precision oncology program either by biopsy or surgery of primary or metastatic lesions. Eligibility criteria and workflow as well as clinical data acquisition are described elsewhere (8,9). The study was conducted in accordance with the Declaration of Helsinki. All patients provided written informed consent of tissue collection as approved by the Universities Ethics Review Boards (University Hospital Heidelberg 323/2004, MASTER Dresden EK431102015, MASTER Heidelberg S-206/2011, MASTER Essen 13-5611-BO).

### Primary cell isolation and generation of patient-derived cancer models (PCM)

For generation of PCMs, cells were freshly isolated from patient material or patient-derived xenografts (PDX) and dissected into 1-2 mm pieces until single-cell suspension or small cell aggregates were obtained using the Miltenyi human Tumor Dissociation kit, (Miltenyi, #130-095-929). Dissociated cells were filtered through a 40 µm Cell Strainer (NeoLab, #352340), and **(i)** cryopreserved in freezing solution (55% Advanced-Dulbecco’s Modified Eagle Medium (DMEM/F12; Life Technologies, #12634010), 30% Fetal Calf Serum (FCS; PAN Biotech, #P40-37500) and 15% DMSO (Sigma-Aldrich, #D4540)), **(ii)** seeded for immediate drug testing as short-term (ST-) PCM, or **(iii)** cultured as long-term (LT-) PCM. Additional PCMs were in-house generated as previously described (32–34).

### Cell culture

Organoids were cultivated in growth factor reduced Matrigel (Corning, #356231), at 2000-6000 cells/well under entity-specific primary culture conditions (35,36). Spheroids and semi-adherent PCMs were cultured as described (32–34) in ultra-low attachment flasks (Corning, #4616 (T25)/#3814 (T75)) or tissue culture-treated flasks (Fisher Scientific, #12034917 (25cm^2^)/#10364131 (75cm^2^)/#10246131 (175cm^2^)), respectively. Long-term PCMs were defined as successfully established when they formed viable microtumor-like structure and expanded for at least 10 passages after initial tumor processing (37). Melanoma cell lines WM3734 and Malme-3M (RRID:CVCL_1438) were cultured in Roswell Park Memorial Institute (RPMI) 1640 (Life Technologies, #21875091) medium with 10% FCS. Pancreatic cancer cell lines NCI-H23 (RRID:CVCL_1547) and Mia-Paca2 were cultured in DMEM (Life Technologies, #11995-065) supplemented with 2.5% Horse Serum (Life Technologies, #26050-070) and 2 mM GlutaMAX (Life Technologies, #35050061), or in RPMI 1640 medium with 10% FCS and 1% PenStrep (Life Technologies, #15140122), respectively.

Isogenic cell lines were generated as previously described (38). Non-malignant pancreatic duct epithelial cell line H6C7 and its isogenic KRAS G12V derivative were cultured in Keratinocyte-SFM Medium (Life Technologies, #17005075) supplemented with L-glutamine, epidermal growth factor (EGF; 5 ng/mL), and bovine pituitary extract (BPE; 50 µg/mL). Non-malignant breast epithelial cell line MCF10A EV (p53 WT) (RRID:CVCL_0598) and the isogenic MCF10A EGFR L858R or KRAS G12D lines were maintained in DMEM/F12 with 5% HEPES buffer (Sigma, #H0887-100ML), 5% Horse Serum, human EGF (20 ng/mL; R&D, #236-EG-01M), hydrocortisone (0.5 mg/mL, Sigma, #H0888-5G), cholera toxin (100 ng/mL, Sigma, #C8052-.5MG), insulin (10 µg/mL, Sigma, #6634-50MG), and PenStrep (10 U/mL). Human embryonic kidney cell line HEK293T (RRID:CVCL_0063) was cultured in IMDM (Life Technologies, # 12440061), including 10% FCS. All long-term PCMs were subjected to Multiplex short tandem repeat (STR)-based authentication (Multiplexion, Heidelberg, Germany) to confirm unique SNP profiles and underwent routine testing for mycoplasma, virus and inter-sample cross-contamination(39). All cell lines were maintained in tissue culture-treated flasks at 37°C and 5% CO2.

### Laboratory Animals and Xenotransplantation

Male and female immunodeficient NOD.Cg-Prkdc^scid^Il2rg^tm1Wjl^/SzJ (NSG) mice were purchased from The Jackson Laboratory (Bar Habor, ME, USA) and bred in the animal facilities of OncoRay, Dresden and the NCT/UCC Dresden under strict specific pathogen-free conditions. For xenotransplantations, 6 to 52-week-old mice were anesthetized on a 37°C heat pad with 1-2.5% isoflurane in the breathing air and 200 mg/kg of Metamizol (WDT) was administered as painkiller. 0.3×0.3 -0,5cm^3^(surgery) or ≤0.5 cm length (fine-needle biopsy) sized tumor fragments were transplanted subcutaneously. Mice were euthanized by cervical dislocation after xenograft volumes reached a maximum of 1.5 cm^3^ or at defined study endpoints, and formed PDX were collected for DST. All animal experimentation adhered to national regulations and was reviewed and approved by the institutional animal welfare committee led by the responsible animal welfare officer. The experiments were authorized by the state directorate of Saxony, Germany (approval numbers: TVV49/2018, TVV39/2023).

### Compounds for Drug Sensitivity Testing

All drugs used for *ex vivo* drug sensitivity testing, including their class, targets, lot numbers and suppliers are listed in the supplement (Suppl. Table 2). Compounds were dissolved in DMSO (Sigma-Aldrich, #D4540) or Water, respectively, at 1-190 mM (mainly 10-80 mM) and stored at -80°C.

### Drug Sensitivity Testing and Metabolic Activity Assay

For drug testing, PCM from all sources were dissociated into single-cell suspensions and seeded as short-term cultures on drug test plates. Dissociation was performed by mechanical dissociation, TrypLE treatment (Life Technologies, #12604013), Trypsin-EDTA 0.05% (Thermo Fisher Scientific, #25300054), or Trypsin-EDTA 0.25% (Thermo Fisher Scientific, #25200056), or Accumax (Merck Millipore, #SCR006) at 37°C. Organoid cultures were processed by mechanical dissociation and Matrigel removal using Cell Recovery Solution (Corning, #354253) on ice for 45 min. Cells derived from patient tumors were first allowed to equilibrate for 0-72 hours. Cells were further dissociated, counted and seeded at densities ranging from 50 to 2000 cells/well in screening medium (DMEM/F12; 10 mM HEPES; 1x MEM non-essential amino acids solution (LifeTechnologies; #11140035); 3.25 mM L-glutamine, 1x B27 supplement, without vitamin A (LifeTechnologies; #12587010); 2 ng/mL heparin (Sigma; #H3149-50KU); 10 U/mL Pen/Strep; 20 ng/mL hEGF, 10 ng/mL PDGF-AA (PeproTech; #100-13A)) onto the ready-to-use plates. For 20-point concentration-response testing, cells were seeded and acclimatized for 24 hours before drug treatment in dilutions of 20 distinct concentrations (n=3) in quadruplicate wells. Miniaturized assays used 5 concentrations in duplicates. 72 hours post seeding ATP levels were quantified using Cell Titer Glo 2.0 (Promega, #G9242) or ATPlite 1step Luminescence Assay (Perkin Elmer, #6016731) according to manufacturer’s instructions. Luminescence signals were measured using Tecan Spark Cyto Reader (TECAN). PCM were tested in one to four biological replicates.

### Pre-Spotted Drug Plates

*Ex vivo* sensitivities to 87 drugs were measured on up to four pre-spotted 384-well ultra-low attachment assay plates (Corning, #3830). Each plate contains a unique set of up to 28 pre-spotted clinically relevant drugs or their in vitro active substances that target key signaling pathways in oncology. Drugs were enriched for FDA-approved or targeted agents relevant to biomarker-driven therapy decisions in rare and molecularly heterogeneous cancers. Using the TECAN D300e Digital Dispenser (TECAN), drugs, DMSO/Water Tween (negative control) benzethonium chloride and staurosporine (positive controls) were added randomized in a 200fold higher concentration than the desired final concentration to a 384-well ultra-low attachment V-shaped stock plate (Thermo Scienitific, #4309). Drugs were tested in duplicates at five different concentrations spanning the compound’s Cmax (when available). Drugs were copied from a stock plate to 384-well ultra-low attachment plates using Mosquito Liquid Handler (TTP LabTech Ltd). Stock plates were sealed with a silver seal (Greiner Bio-One SILVERseal™, #676090). All plates were stored in an oxygen- and moisture-free environment (Dundee StoragePod, Roylan Developments, Fetcham, Leatherhead, UK) at room temperature (DMSO) or -80°C (Water-Tween) until further use. After adding cell suspension, the final DMSO/Water-Tween concentration was 0.5%.and 0.5%, respectively.

### Systematic Drug Sensitivity Identification

Raw luminescence data from all drug-treated and control wells were normalized to plate-specific positive and negative controls to calculate percent inhibition/well. To ensure data quality before downstream analysis, basic quality control was applied at the plate level. Outliers among controls were identified using the Median Absolute Deviation (MAD), where a data point was flagged as an outlier if its absolute deviation from the median exceeded 1.5x MAD (40). As number of control wells per sample varied (range 4-14), outlier removal followed a proportional rule: two outliers were permitted for 4 wells and up to 7 for 14 wells (50%). This approach balanced data integrity with the small replicate number, acknowledging that even a single technical deviation can markedly bias sensitivity estimation (41). Screen performance was evaluated using the QCN-mod module of the iTReX platform (17,18), which provides standardized metrics for high-throughput screening quality control. Quality was passed when there was a distinct signal separation between positive and negative control, which was determined using a zprime_R≥0 (17,18). Only experiments fulfilling the quality control (QC) threshold (zprime_R≥0) were included in downstream analyses. Individual PCM drug sensitivities were then calculated using the iTreX R/Shiny implementation (https://github.com/iTReX-Shiny/iTReX), employing the adjusted asymmetric Drug Sensitivity Score (DSS_asym,adj_) (18). For simplicity, DSS_asym,adj_ was referred to as “DSS” throughout the text. For all miniaturized assays (technical validation, mutation-specific testing, PCM screens), QC thresholds (zprime_R≥0) and the 5-point concentration design were applied, while early exploratory experiments used the original 20-point format.

For patient drug response data, drugs were classified as responders if DSS≥10 and the following quality and sensitivity criteria were met: (i) goodness of fit GOF (R^2^≥0.8) for the inhibition curve (ii) dynamic range ΔPI5-PI1,defined as the difference between maximum and minimum inhibition across tested concentrations, which needed to exceed 75% (ΔPI5-PI1≥75%), and (iii) IC50 below the *in vivo* Cmax of the corresponding drug **(Suppl. Fig. 4)** (17,18). To contextualize each PCM’s response within the total cohort, DSS quantiles were computed for each drug, representing the proportion of models showing lower sensitivity than that of the sample of interest (17).

Dose-response Curve Generation and Parameter Extraction: IC50 graphs were generated by non-linear regression ([Inhibitor] vs. normalized response) after normalizing raw luminescence to plate-specific positive (∼0% cell viability) and negative (∼100% cell viability) controls. IC50 was defined as the drug concentration reducing viability by 50%.

Statistical Analysis of Responders vs. Non-Responder Groups: Sensitivity criteria from iTReX output were used to stratify samples into responders (DSS≥10 and IC50<Cmax) and non-responders. Normalized readout values were multiplied by 100 to represent % viability. Dose-response curves for each response group were generated using the nplr package (version 3.1-0), with the interquartile range shown as ribbon around the fitted curve.

*In silico* Simulations for Low-Input Feasibility Assessment: To systematically assess miniaturization effects, we performed *in silico* simulations based on experimentally acquired high-resolution dose-response curves from PCM77 cells treated with vorinostat and simulated reducing the number of drug concentrations in the dilution series in silico, assessing the impact on key pharmacological readouts. For the concentration format miniaturization, we started from the original 20-concentration format and reduced the number of drug concentrations evenly to obtain 10-, 5- and 3-point concentration series per drug and assessed preservation of original curve shape, concentration spacing and noise structure to allow unbiased comparison between formats.

### Bulk Whole Exome/Genome Sequencing and Transcriptome Analysis of MASTER Patients

Tumor tissues from patients enrolled in the precision oncology program MASTER were drug sensitivity tested and submitted to whole-exome or whole-genome sequencing as described previously (9). Data processing was performed using the DKFZ one-touch pipeline (OTP) (42).

Sequencing and Variant Calling: Whole genome and exome sequencing (WGS and WES) reads were mapped to the human reference genome (hs37d5) and to the genome of Enterobacteria phage phiX174 using BWA mem (version 0.7.15, RRID:SCR_010910). Somatic single nucleotide variants (SNVs) were identified from matched tumor/control sample pairs using SAMtools (version 0.1.19, RRID:SCR_002105) mpileup and bcftools-based pipeline with heuristic filtering (42). SNVs were annotated with ANNOVAR (version Nov. 2014, RRID:SCR_012821) using GENECODE (release 19). SNVs with exonic classification (nonsynonymous, stopgain, stoploss) or ANNOVAR splicing function were included in downstream analyses and visualization. Short insertions and deletions (indels) were detected using Platypus (version 0.8.1, RRID:SCR_005389) and annotated with ANNOVAR (version Feb. 2016). Indels with exonic classification (stopgain or stoploss, frameshift deletion, frameshift insertion, nonframeshift deletion, nonframeshift insertion, or splicing) were visualized in the oncoprint.

Copy Number and Structural Alterations: Copy number alterations (CNA) were inferred using ACEseq (version 5.1.0, for WGS) and CNVkit (version 2.1.0, RRID:SCR_021917, for WES) (42). Optimal tumor cell content and ploidy states were estimated algorithmically and manually reviewed to select the most plausible solution. An algorithm estimated optimal tumor cell content and ploidy combinations, which were visually reviewed to select the most plausible solution. CNAs showing homozygous deletions or amplifications were visualized in the oncoprint.

Microsatellite instability (MSI) status was determined using MSIsensor Pro (version 1.2.0, RRID:SCR_006418) (43). The MSIsensor scan command was applied to the 1000 Genomes reference (RRID:SCR_008801), yielding a reference set of 33,386,244 homopolymer and microsatellite loci.

Transcriptome Analysis and Gene Fusions: RNAseq reads were aligned to the same hs37d5 reference genome using STAR (version 2.5.1b, RRID:SCR_004463) (44). Gene fusions were detected with the Arriba pipeline (version 2.4.0, RRID:SCR_025854) (45). Only high confidence fusion events affecting exon, intron, splice-site, UTR, 3’UTR, or 5’UTR sites were retained for visualization.

Integrated Data Visualization and Expression Analysis: Tumor suppressors and oncogenes, as defined by the COSMIC Cancer Gene Census (Version 05.10.2023, RRID:SCR_002260), were displayed in oncoprints generated with ComplexHeatmap (version 2.22.0, RRID:SCR_017270) (46) if mutated in ≥10% of patients.

### DNA sequencing and Bioinformatic Analysis of LT-PCMs

DNA was extracted from PCMs using the DNeasy Blood & Tissue Kit (Qiagen, Hilden, Germany) according to the manufacturer’s instructions, and processed as described (47). Briefly, 120 ng of total DNA were used for library preparation with the TruSight Oncology 500 Panel Kit (TSO500, Illumina), and the barcoded libraries were sequenced in paired-end (PE) 150bp mode on the NextSeq500/550 systems (Illumina). DNA from blood was processed using the same protocol; germline samples were pooled in equimolar ratios to distinguish between germline and somatic variants. Variant calling was performed using the Illumina TruSight Oncology 500 v2.2 Local App. Resulting small variants (SNVs and indels) were quality-filtered according to minimum variant coverage of 75X and minimum allele frequency of 5%. Information on population frequency (gnomAD_exomes.r2.1.1, RRID:SCR_014964), clinical relevance (ClinVarRRID:SCR_006169), and somatic cancer relevance (COSMIC) was annotated using SnpSift(RRID:SCR_015624), based on the GRCh37.p13 RefSeq reference; variant effects were predicted using SnpEff(RRID:SCR_005191), with the options -no-intergenic, -no-upstream, and -no-downstream applied to exclude non-coding flanking variants. To exclude common variants, a maximum gnomAD allele frequency of 5% was applied. Intron variants without any ClinVar annotation indicating clinical relevance were excluded (benign, likely_benign, benign/likely_benign). If retrieved in the pooled normal sample, the germline variant allele frequency was annotated to the variants of the PCM samples and used to determine somatic relevance (germline variant allele frequency <= 5%).

### Mutational Profiling of Cell Lines

Mutation data for cell lines WM3734, Malme-3M, Mia PaCa-2, and NCI-H23 were obtained from publicly available resources, including the Cancer Cell Line Encyclopedia (CCLE, RRID:SCR_013836) and the COSMIC Cell Lines Project. These datasets provided annotated information on somatic single nucleotide variants (SNVs), small insertions and deletions (indels), and copy number alterations. The mutation profiles were used to confirm the reported oncogenic driver alterations relevant for interpreting mutation-specific drug responses.

### Proliferation Class Assignment

To compare for PCM-type-specific differences in drug sensitivity, ST-PCMs >80% TCC were selected and compared to LT-PCMs. ATP readouts from the drug screening assay normalized to the seeded cell number per well were used as a surrogate for proliferative activity. An unsupervised k-means clustering approach was performed as previously described (48). The resulting cluster centers (i.e., mean ATP readout for each cluster) were extracted and sorted in ascending order. For downstream analyses, PCMs were assigned to the proliferation class corresponding to the cluster with the closest center and only full drug screens (plates 1-3) of PCMs were considered to prevent drug-or drug-class bias. To evaluate differences in normalized ATP readouts, the Wilcoxon rank-sum test was performed. Multiple pairwise comparisons were conducted within drug classes, stratified by proliferation class and PCM type. To control for FDR, Benjamini-Hochberg correction was applied, and adjusted p-values <0.05 were considered statistically significant. Differences in DSS values between proliferation classes were assessed analogously using Wilcoxon rank-sum tests with the same FDR correction strategy.

### Clinical Data Acquisition

To correlate *in vitro* predicted drug sensitivity with clinical outcome, we assessed the first line of therapy administered after the time point of tissue sampling for DST and then compared all therapy lines after sampling. Clinical benefit was primarily assessed based on the best radiologic or clinical response under the tested drug, which was frequently administered as part of a combination therapy regimen. Response evaluation was performed in the palliative or neoadjuvant setting, primarily by CT imaging, provided that no confounding therapies (e.g. radiotherapy or surgical resections) were administered concurrently. Whenever feasible, response evaluation followed RECIST 1.1 criteria (49), and non-RECIST assessments were based on physician-documented radiologic or clinical benefit. Multidisciplinary molecular tumor board (MTB) discussions were used to support response assessment whenever available.

Complete Response (CR) was defined as absence of any radiologically detectable tumor. Partial response (PR) was defined as measurable reduction in tumor size with residual disease. Stable disease (SD) was defined as absence of significant tumor shrinkage or progression, with no new lesions. Progressive disease (PD) was defined as increase in tumor size or appearance of new lesions. A mixed response (MR) was assigned when simultaneous shrinkage of some lesions and progression of others occurred; MR was counted as clinical benefit, consistent with established precision oncology reporting conventions. For response categorization, complete response (CR), partial response (PR), and mixed response (MR) were grouped as response, whereas progressive disease (PD) was classified as non-response. Stable disease (SD) was analyzed as a separate category.

As a second outcome measure, we calculated a progression-free survival ration (PFSr) for the first therapy after tissue sampling. PFSr was defined as the ratio of progression-free survival under the post-sampling therapy (PFS2) to that under the preceding line of therapy (PFS1), as previously described (50). A PFSr ≥ 1.3 was considered indicative of clinical benefit, in line with prior precision oncology studies, whereas PFSr < 1.0 indicated no benefit.

Statistical analyses were using the Wilcoxon rank-sum test or the Kruskal-Wallis test followed by *post hoc* Dunn’s test for multiple comparisons. P<0.05 was considered statistically significant.

### Data Analysis and Statistical Information

All statistical analyses were conducted with the R software (version 4.3.2, 4.4.1). Depending on data distribution and sample size, appropriate statistical tests were applied, including Spearman’s rank correlation, Wilcoxon rank-sum test (two-sided), one-way ANOVA, Mann-Whitney U test, and the Kruskal-Wallis test followed by post hoc Dunn’s test for multiple group comparisons. For multiple testing, the Benjamini-Hochberg correction was applied to control the false discovery rate (FDR). Data were considered statistically significant when p<0.05. For experiments with n≥3, results are expressed as mean ± SD. Dose-response curves were generated using the drc package (version 3.0-1) with a five-parameter logistic model. QC metrics, sensitivity criteria, and DSS calculations (including zprime_R and IC50, GOF) were generated using the iTReX platform as described above.

## Data Availability

All data generated during this study are available from the corresponding author upon reasonable request. Raw reads are available from EGA (EGAD00001015778: Transcriptome sequencing dataset of rare tumors from the MASTER precision oncology registry, EGAD00001015777: Whole-genome sequencing dataset of rare tumors from the MASTER precision oncology registry, EGAD00001015804: Whole-exome sequencing dataset of rare tumors from the MASTER precision oncology registry) under controlled access.

## Acknowledgments

The authors express profound gratitude to all patients and their families for their invaluable contributions to the DKFZ/NCT/DKTK MASTER program. The authors are grateful to the NCT/DKFZ Sample Processing Laboratory, the DKFZ Next Generation Sequencing Core Facility, and the DKFZ Omics IT and Data Management Core Facility for technical support. We express our gratitude to the Core Unit for Molecular Tumor Diagnostics (CMTD) Dresden for conducting sequencing and bioinformatics services for PCM sequencing. In particular, we are grateful to Madeleine Rickauer, Tanja Scholte, as well as the team of the Patient-derived Tumor Model Unit at NCT/UCC Dresden for technical support and coordination. We thank the management and technicians of the animal facility in Oncoray (Dresden) and at NCT/UCC Dresden for their exceptional support. Additionally, we would like to acknowledge the study nurses involved in the monitoring, organization and logistics of tumor samples from patients enrolled in MASTER and Katja Beck and Maria-Veronica Teleanu for administrative support. This work was supported by the NCT Molecular Precision Oncology Program, DKFZ-Heidelberg Center for Personalized Oncology [grant number H021], and DKTK Joint Funding Program and by grants from the BMBF, i.e. the HEROES-AYA (Heterogeneity, Evolution, and Resistance in Oncogenic Fusion Gene-Expressing Sarcomas Affecting Adolescents and Young Adults) consortium within the National Decade Against Cancer of the German Federal Ministry of Education and Research (01KD2207). LM was supported by the German Research Foundation (DFG) through a Walter Benjamin Fellowship (grant number 551924272 and 574708936). M.W. was supported in part by the MeDDrive and ForwarDD Clinician Scientist Programs of the Faculty of Medicine at the Dresden University of Technology. Workflow and timeline figures were generated with Biorender.com. During the preparation of this work, the authors used ChatGPT (OpenAI) to assist with text formulation, specifically to improve wording, text flow, and grammar. All content was carefully reviewed and edited by the authors to ensure scientific accuracy and consistency. The authors take full responsibility for the final version of the manuscript.

## Author Contributions

**Conceptualization:** C.R.B., H.G.

**Study supervision:** C.R.B.

**Data collection & Analysis:** Ja.P., Z.I.C., F.G.T., J.S., L.K.S.F., A.J., A.K., C.D., P.P.T., S.H., S.S., R.P., Je.P., M.G.P.S., E.Y., I.W., M.H., D.R.

**Resources (Samples, Clinical Data & Funding):** Do.H., M.W., I.M., D.E.S., B.S., Da.H., M.S., V.V., C.He., S.K., J.K., O.W., I.O., P.H., L.M., I.K., S.M.P., J.W., K.D.S., D.R., C.H., C.R.B., H.G., S.F., I.M., J.W., C.S.

**Writing – Original Draft & Review:** Ja.P., C.R.B., F.G.T., D.H, L.K.S.F., A.J.

**Writing – Review & Editing:** C.R.B., Ja.P., Z.I.C., F.G.T., J.S., Do.H, L.K.S.F., A.J., A.K., C.D., Je.P., P.P.T., S.H., M.W., M.G.P.S., E.Y., I.W., M.H., I.M., S.S., H.P., D.E.S., B.S., Da.H., C.S., M.S., A.A.W., I.O., V.V., C.He., S.K., P.H., L.M., I.K., S.M.P., J.W., K.D.S., D.R., C.H., S.F., H.G.

**Visualization & Software:** Ja.P., F.G.T., Do.H, L.K.S.F., A.J.

## Abbreviations

3D: Three Dimensional
BztZCl: Benzethoniumchloride
Cmax: Maximum Drug Serum Concentration in vivo
CR: Complete Response
CRC: Colorectal Cancer
CT: Computed Tomography
DMSO: Dimethyl Sulfoxide
DSS: Drug Sensitivity Score
DST: Drug Sensitivity Testing
CI: Confidence Intervals
CUP: Cancer of Unkown Primary
CNV: Copy Number Variants
FDR: False Discovery Rate
GC: Gastric Cancer
GOF: Goodness of Fit
HNC: Head and Neck Carcinoma
IC50: Half-maximal Inhibitory Concentration
Hdels: Homozygous Deletions
HRD: Homologous Recombination
Indels: Small Insertions or Deletions
LC: Lung Carcinoma
LOH: Loss of Heterozygosity
LST: Large-scale State Transitions
LT: Long-Term
MAD: Median Absolute Deviation
MR: Mixed Response
MSI: Microsatellite Instability
N₂: Nitrogen Gas
NSCLC: Non-small Cell Lung Cancer
NET: Neuroendocrine Tumor
NSG: NOD scid gamma
OC: Ovarian Carcinoma
PR: Partial Response
PDAC: Pancreatic Ductal Adenocarcinoma
PCM: Patient-derived Culture Models
PDX: Patient-derived Xenograft
PFS1: Progression-free Survival Time 1
PFS2: Progression-free Survival Time 2
PFSr: Progression-free Survival Ratio
PI1: Percentage Inhibition at 1st drug concentration
PI5: Percentage Inhibition at 5th drug concentration
QC: Quality Control
r: Spearman’s Rank Correlation
RCC: Renal Cell Carcinoma
RNAseq: RNA sequencing
RT: Room Temperature
SARC: Sarcoma
SC: Sensitivity Criteria
sCNAs: Somatic Copy Number Alterations
SD: Stable Disease
SNV: Single Nucleotide Variant
Sph: Spheroid
ST: Short-Term
TCC: Tumor Cell Content
TMB: Tumor Mutational Burden
TSO500: TruSight Oncology 500
WCSS: Within-cluster Sum of Squares

