## Supplemental information for "Systematic functional drug testing in patient-derived models reveals ex vivo sensitivities associated with clinical outcome in rare solid tumors"

#### Supplementary Information Inventory

1. **Supplemental Table 1**: Patient Characteristic Overview of the Tested PCMs.
2. **Supplemental Table 2**: Overview of the used drugs in the DST.
3. **Supplemental Figure 1**: DST Assay Minitaturization and Format Optimization.
4. **Supplemental Figure 2**: Extended Drug Sensitivity Comparison between Short-Term and Long-term PCM.
5. **Supplemental Figure 3**: Landscape of Molecular Alterations Among Patients with Rare Cancers.
6. **Supplemental Figure 4**: Systematic Hit-Identification.
7. **Supplemental Figure 5**: Landscape of Drug Sensitivity Across PCMs of Rare Cancers.

**Supplemental Table 1 Patient Characteristic Overview of the Tested PCMs.**

Baseline demographic and clinical characteristics of patients included in the analysis (n=126).

| | | **Patients (n=126)** |
|---|---|---|
| **Age at Diagnosis (years)** | Mean (SD) | 51 (15.4) |
| | Median (Min, Max) | 53.7 [16.0, 75.2] |
| **Sex** | Female | 54 (42.9%) |
| | Male | 72 (57.1%) |
| **Metastases at Primary Diagnosis** | M0 | 61 (48.4%) |
| | M1 | 65 (51.6%) |
| **Procedure** | Biopsy | 76 (60.3%) |
| | Surgery | 50 (39.7%) |
| **Tumor Sample** | Local Recurrence | 17 (13.5%) |
| | Metastasis | 86 (68.3%) |
| | Primary | 23 (18.3%) |
| **Single Systemic Antineoplastics prior Sampling** | Mean (SD) | 3.28 (2.49) |
| | Median (Min, Max) | 3.00 [0, 12.0] |
| **Radiation before tissue sampling** | No | 85 (67.5%) |
| | Yes | 41 (32.5%) |

**Supplemental Table 2 Overview of the used drugs in the DST.**

| | | | | | Drug Concentration | | | | |
|---|---|---|---|---|---|---|---|---|---|
| Plate | Inhibitor Name | Class | Target | Cmax | V high | V low | Supplier | Item Number | Lot Number |
| 1 | Afatinib | Protein Tyrosine Kinase inhibitor | EGFR (wt & mut), HER2, HER4 | 0.052 | 20 | 0.001 | Selleck Chemicals | S1011-5mg | S101111 |
| 1 | Alectinib | Protein Tyrosine Kinase inhibitor | ALK, RET | 1.38 | 20 | 0.01 | MCE/Hölzel Diagnostika | HY-13011-5mg | 283624 |
| 1 | Axitinib | Protein Tyrosine Kinase inhibitor | VEGFR1-3, c-KIT, PDGFR, | 0.16 | 50 | 0.001 | Focus Biomolecules/Tebu Bio | 21910-2118-10mg | X106716 |
| 1 | Cabozantinib | Protein Tyrosine Kinase inhibitor | VEGFR, c-MET, RET, KIT, FLT3, TIE2, AXL | 4.61 | 25 | 0.01 | Selleck Chemicals/Hoelzel Diagnostika | S1119 | S111912 |
| 1 | Cobimetinib | MAPK inhibitor | MEK1/2 | 0.51 | 100 | 0.001 | Selleck Chemicals/Hoelzel Diagnostika | S8041-5 | S804103 |
| 1 | Copanlisib | PI3K/Akt/mTOR inhibitor | PI3K α/δ, PI3K β/γ | 0.96 | 2.5 | 0.0001 | Hoelzel Diagnostika | TMO-T6322-2mg | 120379 |
| 1 | Crizotinib | Protein Tyrosine Kinase inhibitor | ALK, c-MET, ROS1, RON, AXL | 0.91 | 100 | 0.001 | Selleck Chemicals/Hoelzel Diagnostika | S1068-10 | S106817 |
| 1 | Entrectinib | Protein Tyrosine Kinase inhibitor | ALK, ROS1, Trk A/B/C, JAK2 | 3.13 | 100 | 0.001 | Selleck Chemicals/Biozol | S7998-25 | S799801 |
| 1 | Everolimus | PI3K/Akt/mTOR inhibitor | mTORC1, FKBP-12 binding | 0.06 | 10 | 0.001 | Selleck Chemicals/Hoelzel Diagnostika | S1120-10 | S112026 |

| | | | | | Drug Concentration | | | | |
|---|---|---|---|---|---|---|---|---|---|
| Plate | Inhibitor Name | Class | Target | Cmax | V high | V low | Supplier | Item Number | Lot Number |
| 1 | Imatinib | Protein Tyrosine Kinase inhibitor | Bcr-ABL, c-KIT, PDGFR, ABL1, DDR1 | 7.50 | 100 | 0.01 | Selleck Chemicals/Hoelzel Diagnostika | S1026-100 | S102607 |
| 1 | Ipatasertib | PI3K/Akt/mTOR inhibitor | pan-Akt | 1.06 | 100 | 0.001 | Selleck Chemicals/Hoelzel Diagnostika | S2808-10 | S280805 |
| 1 | Lapatinib | Protein Tyrosine Kinase inhibitor | EGFR, HER2 | 4.18 | 10 | 0.001 | Selleck Chemicals/Biozol | S2111-25 | S2111107 |
| 1 | Larotrectinib | Protein Tyrosine Kinase inhibitor | Trk A/B/C | 1.84 | 100 | 0.001 | Selleck Chemicals/Hoelzel Diagnostika | S7960-25 | S796002 |
| 1 | Nintedanib | Protein Tyrosine Kinase inhibitor | FGFR1-3, PDGFR, VEGFR1-3, FLT3, RET | 0.0004 | 0.001 | 0.0000065 | MCE/Hölzel Diagnostika | HY-50904-10mM | 103323 |
| 1 | Olaparib | PARP inhibitor | PARP | 13.12 | 100 | 0.01 | Selleck Chemicals/Biozol | S1060-10 | S106023 |
| 1 | Palbociclib | Cell Cycle inhibitor | CDK4/6 | 0.10 | 10 | 0.001 | Selleck Chemicals/Biozol | S1116-5 | S111613 |
| 1 | Panobinostat | Epigenetic modifier | HDAC | 0.08 | 1 | 0.0001 | Selleck Chemicals/Biozol | S1030-10 | S103013 |
| 1 | Pazopanib | Protein Tyrosine Kinase inhibitor | c-KIT, PDGFR, VEGFR1-3, FGFR, LCK, c-RAF, B-RAF, c-Fms/CSF1R | 133.00 | 100 | 0.001 | Selleck Chemicals/Hoelzel Diagnostika | S3012-25 | S301206 |
| 1 | Regorafenib | Protein Tyrosine Kinase inhibitor | VEGFR 1-3, PDGFR, FGFR, c-RET, c-KIT, RAF-1, TIE2 | 8.08 | 50 | 0.01 | Selleck Chemicals/Hoelzel Diagnostika | S1178-10 | S117805 |

| | | | | | Drug Concentration | | | | |
|---|---|---|---|---|---|---|---|---|---|
| **Plate** | **Inhibitor Name** | **Class** | **Target** | **Cmax** | **V high** | **V low** | **Supplier** | **Item Number** | **Lot Number** |
| 1 | Ruxolitinib | JAK/STAT inhibitor | JAK1/2 JAK3, TYK2 | 1.09 | 100 | 0.001 | Selleck Chemicals/Hoelzel Diagnostika | S5243-25 | S524305 |
| 1 | Selpercatinib | Protein Tyrosine Kinase inhibitor | c-RET | 0.01 | 1 | 0.0001 | Selleck Chemicals | S8781 | S878103 |
| 1 | Sorafenib | Protein Tyrosine Kinase inhibitor | RAF-1, VEGFR1-3, PDGFR, BRAF, FLT3, c-KIT, RET | 20.10 | 200 | 0.01 | Selleck Chemicals/Hoelzel Diagnostika | S7397-20 | S739707 |
| 1 | Sunitinib | Protein Tyrosine Kinase inhibitor | PDGFR, c-KIT, VEGFR1-3, FLT3, RET, CSF-1R | 0.18 | 50 | 0.001 | Selleck Chemicals/Hoelzel Diagnostika | S7781-50 | S778105 |
| 1 | Talazoparib | PARP inhibitor | PARP1-2 | 0.026 | 10 | 0.001 | Selleck Chemicals/Hoelzel Diagnostika | S7048-10 | S704808 |
| 1 | Tazemetostat | Epigenetic modifier | EZH2 inhibitor, EZH1 | 1.45 | 50 | 0.01 | Selleck Chemicals/Hölzel | S7128-10 | S712811 |
| 1 | Trabectedin | Chemotherapeutics | DNA binding | 0.002 | 0.02 | 0.000002 | Pharma Mar S.A. | PZN 05704993 | 23164 |
| 1 | Venetoclax | Apoptosis modulator | Bcl-2 | 2.41 | 50 | 0.01 | Selleck Chemicals/Hoelzel Diagnostika | SEL-S8048-50MG | S804820 |
| 1 | Vismodegib | Hedgehog Pathway inhibitor | Hedgehog | 33.90 | 100 | 0.001 | Hoelzel Diagnostika | S1082-10 | S108210 |
| 2 | (+)-JQ-1 | Epigenetic modifier | pan-BET | N/A | 50 | 0.001 | MCE/Biozol | HY-13030-10mg | 84732 |

| | | | | | Drug Concentration | | | | |
|---|---|---|---|---|---|---|---|---|---|
| Plate | Inhibitor Name | Class | Target | Cmax | V high | V low | Supplier | Item Number | Lot Number |
| 2 | Alpelisib | PI3K/Akt/mTOR inhibitor | PI3K-α | 5.6 | 100 | 0.001 | MCE/Biozol | HY-15244-10mg | 149875 |
| 2 | AMG-900 | Protein Ser/Thr Kinase inhibitor | pan-Aurora | 3.3 | 25 | 0.001 | Selleck Chemicals/Hoelzel Diagnostika | S2719-10 | S271901 |
| 2 | Pasireotide | Somatostatin analogue | Somatostatin Receptor type 1, 2, 3, (4, lower affinity), 5 | 0.00533 | 0.025 | 0.00001 | Selleck Chemicals/Biozol | SEL-P1127-1MG | P112701 |
| 2 | Ceritinib | Protein Tyrosine Kinase inhibitor | ALK, IR, IGF1R,ROS1, FLT3, STK22D | 1.20758 | 10 | 0.001 | MCE/Biotrend | HY-15656-5mg | 19216 |
| 2 | Chloroquine | Chemotherapeutics | Autophagy, ATM/ATR | 0.725 | 100 | 0.001 | Selleck Chemicals/Hoelzel Diagnostika | S4157 | S415703 |
| 2 | Dactolisib | PI3K/Akt/mTOR inhibitor | Pan-PI3K, mTOR1-2, (p70S6K) | 0.07 | 0.7 | 0.0001 | Selleck Chemicals/Hoelzel Diagnostika | S1009-50 | S100915 |
| 2 | Dasatinib | Protein Tyrosine Kinase inhibitor | Bcr-Abl, Src, DDR2, c-KIT, PDGFR, EPHA2 | 0.264 | 100 | 0.001 | Cayman/Biomol | Cay11498-10 | 0621991-49 |
| 2 | Decitabine | Chemotherapeutics | DNA Methyltransferase | 0.323 | 100 | 0.001 | Selleck Chemicals/Biozol | SEL-S1200-10MG | S120011 |
| 2 | Dinaciclib | Cell Cycle inhibitor | CDK1, CDK2, CDK5, CDK9 | 0.36 | 1 | 0.0001 | Selleck Chemicals/Biozol | SEL-S2768-10MM/1ML | S276807 |

| | | | | | Drug Concentration | | | | |
|---|---|---|---|---|---|---|---|---|---|
| **Plate** | **Inhibitor Name** | **Class** | **Target** | **Cmax** | **V high** | **V low** | **Supplier** | **Item Number** | **Lot Number** |
| 2 | Eprenetapopt | Apoptosis modulator | mutant p53 (p53 reactivator), Redox | 286.57 | 100 | 0.001 | Selleck Chemicals/Biozol | SEL-S7724-5MG | S772403 |
| 2 | Erdafitinib | Protein Tyrosine Kinase inhibitor | pan-FGFR, c-RET, CSF-1R, PDGFR, FLT4, c-KIT, VEGFR2 | 3.1 | 100 | 0.001 | MCE/Biozol | HY-18708-10mg | 109837 |
| 2 | Erlotinib | Protein Tyrosine Kinase inhibitor | EGFR, HER2 | 3.14692 | 10 | 0.0001 | Selleck Chemicals/Hoelzel Diagnostika | S1023-10 | S102305 |
| 2 | Ganetespib | Proteosome inhibitor | Hsp90 | 13.7 | 10 | 0.0001 | Selleck Chemicals/Biozol | SEL-S1159-5MG | S115902 |
| 2 | Gilteritinib | Protein Tyrosine Kinase inhibitor | FLT3, AXL | 0.67 | 2.5 | 0.0001 | Selleck Chemicals | S7754 | S775403 |
| 2 | Ivosidenib | Antineoplastics | IDH1 | 11.05 | 1 | 0.0001 | Selleck Chemicals | S8206 | S820603 |
| 2 | Navitoclax | Apoptosis modulator | Bcl-2, Bcl-XL, Bcl-w | 2.62 | 75 | 0.01 | MCE/Hölzel Diagnostika | HY-10087-10mg | 122700 |
| 2 | Neratinib | Protein Tyrosine Kinase inhibitor | EGFR1/2, HER2, HER4 | 0.1369 | 5 | 0.0001 | MCE | HY-32721 | 97011 |
| 2 | ODM-207 | Epigenetic modifier | pan-BET | N/A | 100 | 0.001 | Axon Medchem | Axon 3329 | Batch 01 |
| 2 | Ponatinib | Protein Tyrosine Kinase inhibitor | Bcr-Abl and T315I mutation, FGFR1-4, FLT3, PDGFRα, VEGFR2, c-SRC, c-KIT, DDR2 | 0.137 | 10 | 0.0001 | Selleck Chemicals/Biozol | SEL-S1490-10MG | S1490-10 |
| 2 | Selinexor | Apoptosis modulator | XPO1 | 1.534 | 100 | 0.001 | Selleck Chemicals/Biozol | S7252-5 | S725205 |

| | | | | | Drug Concentration | | | | |
|---|---|---|---|---|---|---|---|---|---|
| Plate | Inhibitor Name | Class | Target | Cmax | V high | V low | Supplier | Item Number | Lot Number |
| 2 | Sonidegib | Hedgehog Pathway inhibitor | SMO | 2.12 | 100 | 0.001 | Selleck Chemicals/Hoelzel Diagnostika | S2151-10 | S725205 |
| 2 | Sotorasib | Protein Tyrosine Kinase inhibitor | KRAS G12C | 13.3784 | 100 | 0.001 | Selleck Chemicals/Biozol | S8955-5 | S895503 |
| 2 | Taselisib | PI3K/Akt/mTOR inhibitor | PI3K α/δ/γ, PI3K β | 0.143 | 100 | 0.001 | MCE/Biozol | HY-13898-10mg | 10696 |
| 2 | Tipifarnib | Metabolism modifier | Farnesyltransferase, H-Ras (activated) | 0.817 | 50 | 0.001 | MCE/Hölzel Diagnostika | HY-10502-10mg | 21104 |
| 2 | Tivantinib | Protein Tyrosine Kinase inhibitor | c-MET | 0.20655 | 100 | 0.001 | Selleck Chemicals/Hölzel | S2753-10 | S275304 |
| 2 | Trametinib | MAPK inhibitor | MEK1/2 | 0.036 | 1 | 0.0001 | Selleck Chemicals/Hoelzel Diagnostika | S2673-5 | S267311 |
| 2 | Vemurafenib | MAPK inhibitor | BRAF V600E | 127 | 50 | 0.01 | Selleckchem | S1267 | S126715 |
| 3 | 5-Fluorouracil | Chemotherapeutics | Pyrimidine antagonist | 1.153 | 10 | 0.001 | Selleck Chemicals/Hoelzel Diagnostika | S1209-100 | S120904 |
| 3 | Abemaciclib | Cell Cycle inhibitor | CDK4/6 | N/A | 1 | 0.0001 | Selleck Chemicals/Hoelzel Diagnostika | S7158-5 | S715803 |
| 3 | BI-3406 | MAPK inhibitor | SOS-1 | N/A | 100 | 0.01 | Selleck Chemicals/Hoelzel Diagnostika | S8916-5 | S891602 |
| 3 | Bleomycin | Chemotherapeutics | DNA synthesis inhibitor | 706 | 150 | 0.01 | Selleck Chemicals/Biozol | S1214-50 | S121421 |

| | | | | | Drug Concentration | | | | |
|---|---|---|---|---|---|---|---|---|---|
| **Plate** | **Inhibitor Name** | **Class** | **Target** | **Cmax** | **V high** | **V low** | **Supplier** | **Item Number** | **Lot Number** |
| 3 | Bortezomib | Proteosome inhibitor | 26s | 0.312 | 3 | 0.00001 | Abmole/Hoelzel biotech | M1686-5mg | M1686-01 |
| 3 | Cyclophosphamide | Chemotherapeutics | DNA Alkylation | 128 | 125 | 0.001 | MCE/Hölzel Diagnostika | HY-117433-10mg | 317261 |
| 3 | Dabrafenib | MAPK inhibitor | BRAF V600E, K, D mutants | 4.86 | 2.5 | 0.0001 | MCE | HY-14660 | 153495 |
| 3 | Doxorubicin | Chemotherapeutics | Topoisomerase II | 6.73 | 1 | 0.0001 | Selleck Chemicals/Biozol | S1208-10 | S120815 |
| 3 | Encorafenib | MAPK inhibitor | BRAF V600E | 12.5 | 10 | 0.0001 | Selleck Chemicals/Hoelzel Diagnostika | S7108-5 | S710801 |
| 3 | Etoposide | Chemotherapeutics | Topoisomerase II | 33.4 | 100 | 0.001 | Cayman/Biomol | Cay12092-25 | 0563006-101 |
| 3 | Gemcitabine | Chemotherapeutics | Further DNA Synthesis inhibition | 89.3 | 100 | 0.001 | Selleck Chemicals/Biozol | S1714-50 | S171404 |
| 3 | Idasanutlin | Apoptosis modulator | MDM2 (restoring p53 activity) | N/A | 20 | 0.001 | Selleck Chemicals/Hoelzel Diagnostika | S7205-25 | S720502 |
| 3 | Palifosfamide | Chemotherapeutics | DNA Alkylation | N/A | 200 | 0.01 | Selleck Chemicals/Hoelzel Diagnostika | S5840-25 | S584001 |
| 3 | JAB-3068 | Protein Tyrosine Phosphatase inhibitor | SHP2 | N/A | 100 | 0.001 | ChemieTek | CT-JAB3068 | #01 |
| 3 | Lenvatinib | Protein Tyrosine Kinase inhibitor | VEGFR1-3, FGFR1-4, PDGFR, c-KIT, c-RET | 0.76139 | 10 | 0.0001 | MCE/Biotrend | HY-10981-5mg | 240149 |

| | | | | | Drug Concentration | | | | |
|---|---|---|---|---|---|---|---|---|---|
| **Plate** | **Inhibitor Name** | **Class** | **Target** | **Cmax** | **V high** | **V low** | **Supplier** | **Item Number** | **Lot Number** |
| 3 | Lorlatinib | Protein Tyrosine Kinase inhibitor | ALK, ROS1, MET | 1.42 | 1 | 0.0001 | Selleck Chemicals/Hoelzel Diagnostika | S7536-5 | S753603 |
| 3 | Osimertinib | Protein Tyrosine Kinase inhibitor | EGFR mutant-selective: EGFRL858R, T790M | 0.126 | 2 | 0.0004 | Selleck Chemicals/Hoelzel Diagnostika | S5078-25 | S507802 |
| 3 | Paclitaxel | Chemotherapeutics | Taxanes (preventing microtubule depolymerization) | 4.3 | 5 | 0.0001 | Selleck Chemicals/Hoelzel Diagnostika | S1150-10 | S115011 |
| 3 | RMC-4630 | Protein Tyrosine Phosphatase inhibitor | SHP2, Impacts various signaling pathways, including RAS/MAPK inhibitor | N/A | 25 | 0.001 | MedChemExpress | HY-141523 | 144764 |
| 3 | Romidepsin | Epigenetics modifier | HDAC | 0.698 | 10 | 0.001 | Selleck Chemicals/Biozol | S3020-5 | S302003 |
| 3 | Selitrectinib | Protein Tyrosine Kinase inhibitor | Trk A/B/C (NTRK gene fusions) | N/A | 0.7 | 0.0001 | MCE/Hölzel Diagnostika | HY-101977-5mg | 41819 |
| 3 | Sitravatinib | Protein Tyrosine Kinase inhibitor | TYRO3, AXL, MER, VEGFR, c-Kit, c-MET, PDGFR | N/A | 5 | 0.001 | Selleck Chemicals/Hoelzel Diagnostika | S8573 | S857301 |
| 3 | SN-38 | Chemotherapeutics | Topoisomerase I | 0.048 | 2 | 0.0004 | Selleck Chemicals/Biozol | SEL-S4908-10MG | S490803 |
| 3 | Temozolomide | Chemotherapeutics | DNA methylation at the O6 position of guanine | 38.6 | 300 | 1 | Selleck Chemicals/Biozol | S1237-25 | S123707 |

| | | | | | Drug Concentration | | | | |
|---|---|---|---|---|---|---|---|---|---|
| Plate | Inhibitor Name | Class | Target | Cmax | V high | V low | Supplier | Item Number | Lot Number |
| 3 | Veliparib | PARP inhibitor | PARP1-2 | N/A | 10 | 0.001 | Santa Cruz | sc-394457A | C1323 |
| 3 | Vincristin | Chemotherapeutics | Inhibiting tubulin polymerization (Vinca alkaloid, Microtubule inhibitor) | 0.007 | 0.01 | 0.00001 | Selleck Chemicals/Hoelzel Diagnostika | S1241-10 | S124108 |
| 3 | Volasertib | Protein Ser/Thr Kinase inhibitor | PLK1, PLK2/3 | 0.79 | 1 | 0.0001 | MCE/Hölzel Diagnostika | HY-12137-5mg | 57030 |
| 3 | Vorinostat | Epigenetics modifier | HDAC | 1.2 | 10 | 0.0001 | Cell Signaling Technology | 12520S | 2 |
| 4 | Cisplatin | Chemotherapeutics | DNA Damage | 14.4 | 25 | 0.1 | Selleck Chemicals | S1166 | S1166 |
| 4 | Carboplatin | Chemotherapeutics | DNA Damage | 135 | 100 | 0.1 | MedChemExpress | HY-17393 | 150608 |
| 4 | Oxaliplatin | Chemotherapeutics | DNA Damage | 4.96 | 12.5 | 0.1 | MedChemExpress | HY-17371 | 25246 |
| Previous drug on plate 2 | Cediranib | Protein Tyrosine Kinase | VEGFR, FLT1/4, c-KIT, PDGFR | 0.1667 | 10 | 0.001 | Selleck Chemicals/Hoelzel Diagnostika | S1017-10 | S101705 |
| Positive Control | Benzethonium Chloride | | | N/A | constant | constant | Selleck Chemicals | S1421-2 | S142107 |
| Positive Control | Staurosporine | | | N/A | 1 | 0.0001 | Selleck Chemicals | S4162 | S416201 |

### Supplemental Legends

**Supplemental Figure 1.**

**DST Assay Minitaturization and Format Optimization.**

**A** Raw value correlation of two technical repeats of three CRC cultures: PCM31, PCM29, and PCM28; three pancreatic cultures: PCM54, PCM55, PCM56; and three sarcoma cultures: PCM77, PCM76, and PCM75 treated with cobimetinib (0.001, 0.1, 1, 10, 100 µM), sorafenib (0.01, 1, 10, 100, 200 µM), and trabectedin (0.000002, 0.00002, 0.0002, 0.002, 0.02) (n=1 biological replicates).

**B-C** Assessment of the relationship between zprime_R (**B**), and relative cell viability (%) (**C**) of primary sarcoma cultures PCM75 with etoposide in five different concentrations (0.001, 0.01, 1, 10, 100 µM) and duplicates (n=1 biological replicate). HEK293T was used as a negative control for etoposide cell viability assessment. Cell viability (%) is relative to the positive (BztCl) and negative (DMSO) control. Respective IC50 values are shown in the legends. The horizontal blue dashed line represents the 50% cell viability in **C.** The red vertical dashed line in **C** marks the Cmax of etoposide. % Viability derived from normalized ATP Readout was plotted against tested drug concentrations using a 5-parameter logistic model.

**D** Interpolated dose-response curves of the average response at overlapping concentrations of manual (n=3 biological replicates) and high-throughput drug screening (n=2 biological replicates) of the sarcoma LT-PCM PCM75 treated with copanlisib in 25 different concentrations for manual DST (0.0000001-5 µM, 2-fold dilution) and quadruplicates and in 5 different concentrations for high-throughput DST (0.0001, 0.001, 0.1, 1.25, and 2.5 µM) and duplicates.

**E-F** Assessment of the impact of storage duration on six plates (plate 2) tested with PCM75 (n=1) after 0, 4, 7, 13, 18, 21 and 27 weeks on **E** relative cell viability (%) after alpelisib inhibition, as well as **F** the z_prime_R.

**G-J** Correlation of DSS across **G-H** two biological repeats of PCM40 (**G**) and PCM95 (**H**) and **I-J** across different sample handling methods, including DST of **I** fresh *vs* frozen of **patient** PCM10 and **J** acclimatized and non-acclimatized samples of PCM10 treated with 84–87 drugs (n=1 biological replicate). The black diagonal represents the correlation line starting at 0. Colored labels indicate the drug class of the specific drug. Spearman's rank correlation coefficient (r) is displayed. Experiments in **G** and **I** were conducted with the previous drug layout, including cisplatin and cediranib and excluding cyclophosphamide and pasireotide. Ave: Average, Neg. Ctrl: Negative control, HTS: High-throughput drug screening, MS: Manual drug screening

**Supplemental Figure 2.**

**Extended Drug Sensitivity Comparison between Short-Term and Long-term PCM.**

**A-L** DSS distribution of samples treated with 84 drugs comprising distinct drug classes. Samples are categorized as either ST-PCM (n=18) with a minimum tumor cell content (TCC) of 80% or LT-PCM (n=53) and further, and stratified according to their proliferation class, determined by an unsupervised k-means clustering approach using ATP Readouts normalized based on seeded cell number. The center line represents the median and the whisker the standard deviation. Significance was determined by the Wilcoxon Rank Sum test with FDR multiple-testing correction. FDR-corrected *p*-values <0.05 were considered significant. ns: not significant; *p<0.05, p<0.01, **p<0.0001. ST-PCM: Short-term patient-derived cancer model, LT-PCM: Long-term patient-derived cancer model

**M-N** Comparison of DSS distribution between LT- (n=53) and ST-PCM (n=18) treated with various drug classes. Boxes illustrate the $25^{th}$- and $75^{th}$-percentiles, respectively, while the center line represents the median. The whiskers extend to the maximum and minimum DSS values observed for the individual drug. The gray vertical dashed line indicates the DSS=10 cutoff.

**Supplemental Figure 3.**

**Landscape of Molecular Alterations Among Patients with Rare Cancers.**

Complex characteristic for each patient's PCM sample presented on an oncoprint. The upper bar plot illustrates the sum of SNVs, indels within exonic sequences per mega base of the coding sequence of the genome, indicating the tumor mutational burden (TMB). Panels directly below show the annotation of (i) entities of the samples, (ii) PCM type, (iii) HRD scores (LOH + LST), (iv) MSI status. Colors in the oncoprint indicate the most frequently observed amplifications (red), hdels (blue), SNVs (purple), indels (orange) and fusions of high confidence (green). Panels below show (i-ii) the sequencing type performed and (iii) tumor cell content (TCC). The bar plots on the right illustrate the percentage of alterations within the gene. hdel: homozygous deletion, indel: short insertions and deletionsSNV: single-nucleotide variant

**Supplemental Figure 4.**

**Systematic Hit-Identification.**

**A** Distribution of cell numbers, categorized by the sample source. Dots are colored according to drug sensitivity screen type. Significance was determined by Mann-Whitney test (two-tailed). ****p<0.0001.

**B** Flowchart of systematic hit-identification: To support patient treatment recommendations in the MTB, drug response was evaluated based on previously described and published quality and sensitivity criteria (QC; SC)(17,18). Readouts were only analyzed if they met the QC criterion zprime_R≥0.5. Additionally, *in vitro* drug responders had to fulfill following QC and SC: (i) goodness of fit ($R^2$, GOF) for the inhibition curve ≥0.8, (ii) DSS≥10, (iii) maximum effect observed between the highest (PI5) and lowest drug concentration (PI1) was at least 75% (ΔPI5-PI1), and (iv) IC50<*in vivo* Cmax of the corresponding drug. Furthermore, DSS values of individual PCMs were compared to the PCM cohort using dot plots and percentiles. DSS values above the $75^{th}$-percentile was considered to indicate an above-average response (TOP25 hit) within the cohort.

**C-F** Identification and categorization of *ex vivo* responders and non-responders after treatment of PCMs with **C** dabrafenib (responders n=2, non-responders n=112), **D** afatinib (responders n=3, non-responders n=156), **E** regorafenib (responders n=15, non-responders n=144) and **F** SN-38 (responders n=30, non-responders n=84). Dose-response curves fitted with a 5-parameter logistic model, serving as a trendline for the responder groups, the interquartile viability across samples within each responder group is indicated as a shaded ribbon around the trendline. Samples are categorized into responder and non-responder groups based on their IC50 values and the corresponding drug's Cmax to define clinical relevance as well as their DSS values. Responders (green): IC50<Cmax & DSS≥10; non-responders (gray): IC50>Cmax & DSS≤10.

**Supplemental Figure 5.**

**Landscape of Drug Sensitivity Across PCMs of Rare Cancers.**

**A** DSS heatmap, annotated with tumor entities, PCM types, and drug classes ordered by hierarchical (Euclidean) clustering of the samples and drugs from all tested samples (n=143) (NA was imputed as -1). **B-D** The percentage of samples in which each drug was classified as a hit was normalized either by the total count of all samples **B** or by the total count of all samples for the respective entity

**C-D**. Colored segments indicate the contribution of each entity to the total hit frequency.

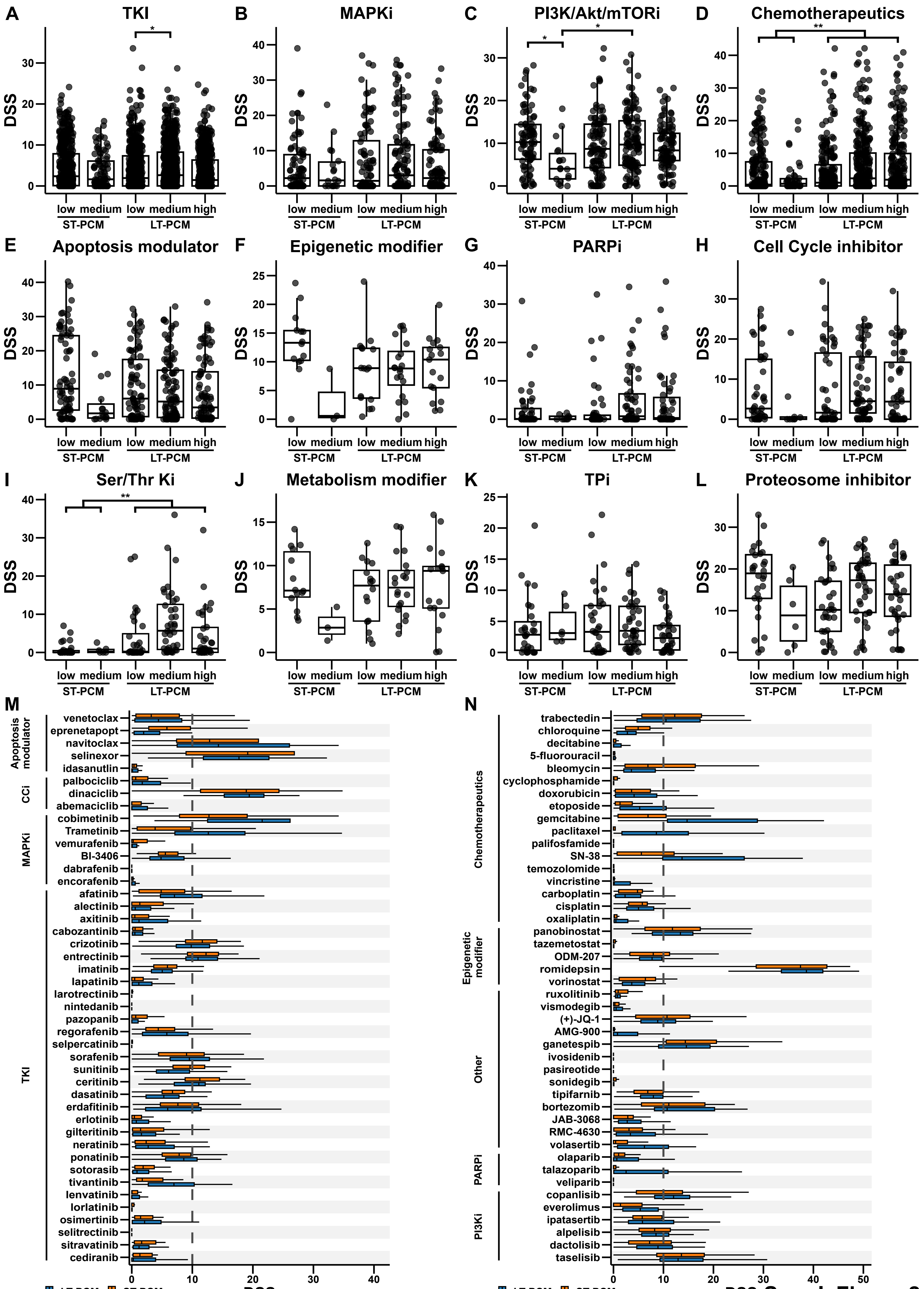


Suppl. Figure 2

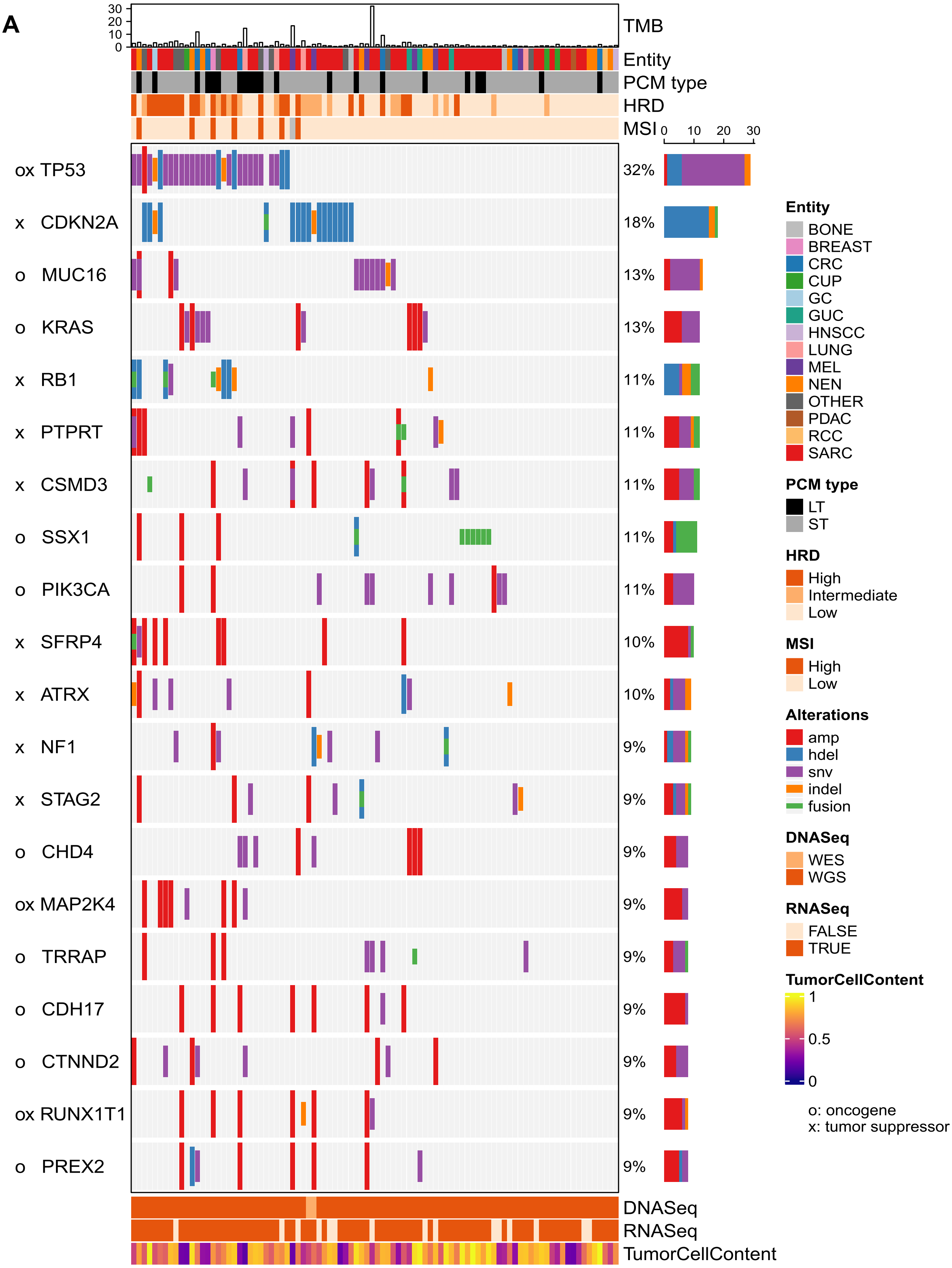


**Suppl. Figure 3**

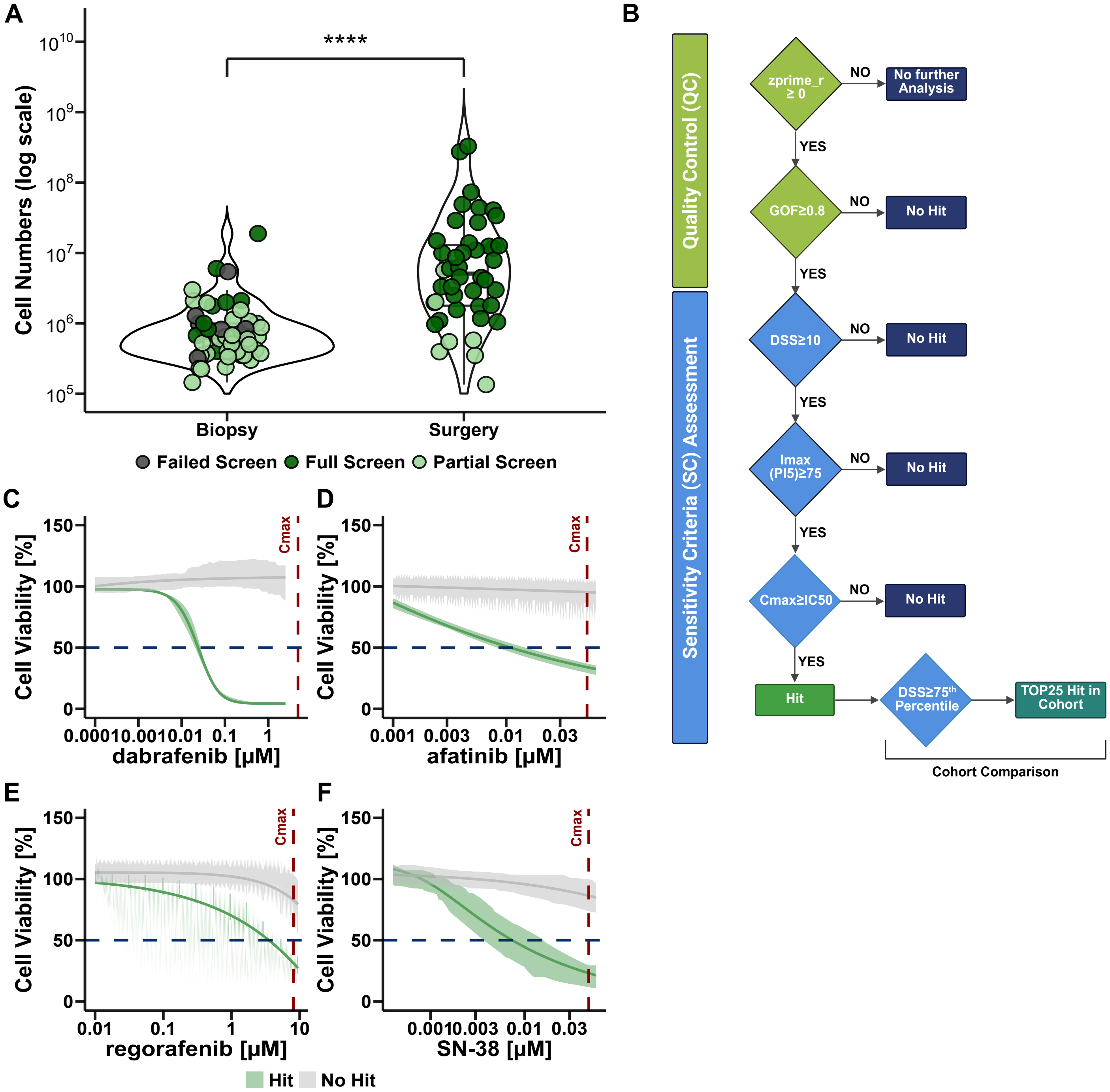


Suppl. Figure 4

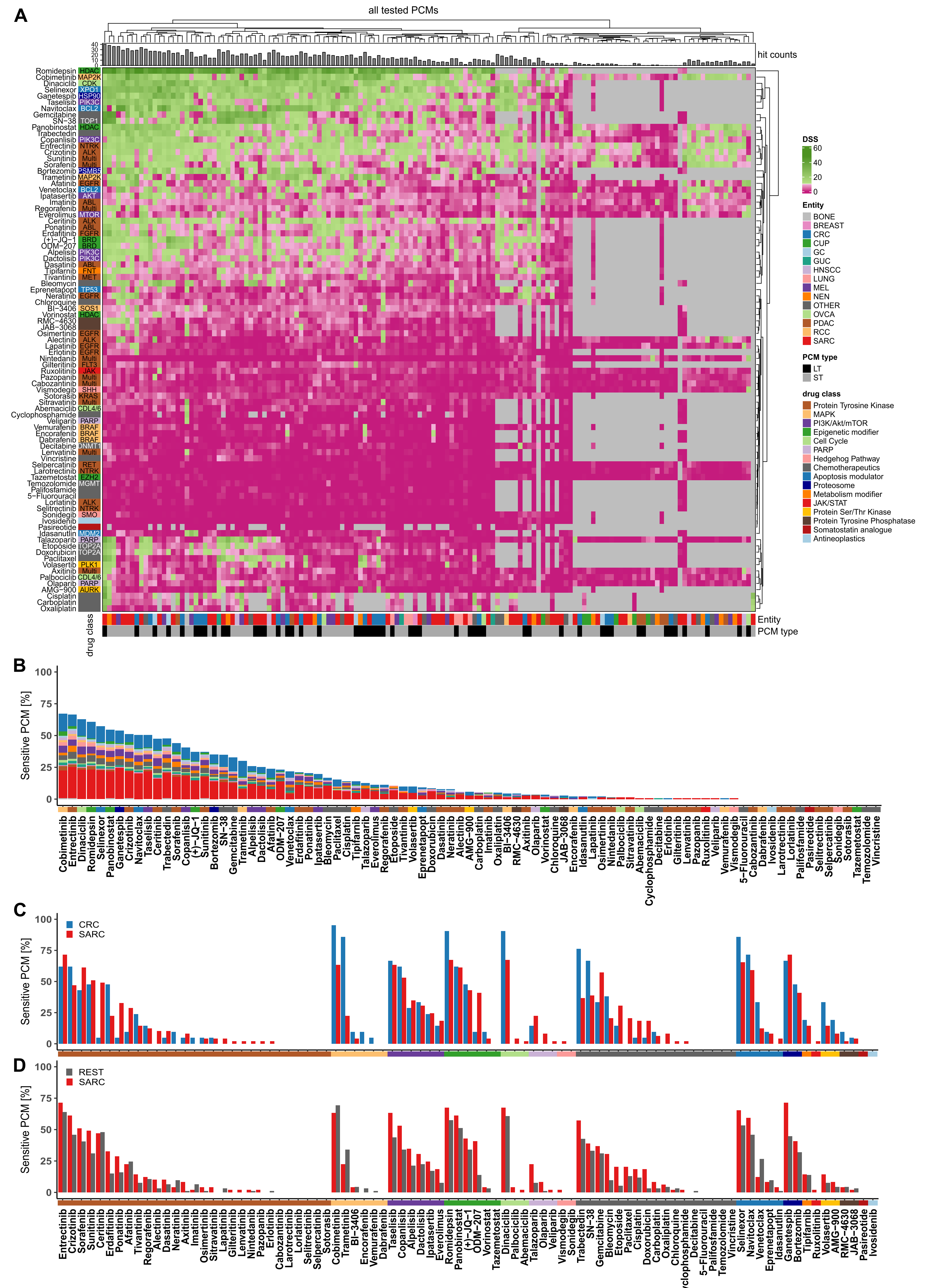


Suppl. Figure 5
